# To impute or not to impute in untargeted metabolomics - that is the compositional question

**DOI:** 10.1101/2024.10.28.620738

**Authors:** Dennis Dimitri Krutkin, Sydney Thomas, Simone Zuffa, Prajit Rajkumar, Rob Knight, Pieter C. Dorrestein, Scott T. Kelley

**Author notes:** Organization: Ometa Labs.

## Abstract

Untargeted metabolomics often produce large datasets with missing values, arising from biological or technical factors, which can undermine statistical analyses and lead to biased biological interpretations. Imputation methods, such as k-Nearest Neighbors (kNN) and Random Forest (RF) regression are commonly used but their effects vary depending on the type of missing data e.g. Missing Completely At Random (MCAR) and Missing Not At Random (MNAR). Here, we determined the impacts of degree and type of missing data on the accuracy of kNN and RF imputation using two datasets: a targeted metabolomic dataset with spiked-in standards and an untargeted metabolomic dataset. We also assessed the effect of compositional data approaches (CoDA), such as the centered log-ratio (CLR) transform, on data interpretation, since these methods are increasingly being used in metabolomics.

Overall, we found that kNN and RF performed more accurately when the proportion of missing data across samples for a metabolic feature was low. However, these imputations could not handle MNAR data and generated wildly inflated values or imputed values where none should exist. Furthermore, we show that the proportion of missing values had a strong impact on the accuracy of imputation which affected the interpretation of the results. Our results suggest extreme caution should be used with imputation even with modestly levels of missing data or when the type of missingness is unknown.

## INTRODUCTION

The exponential growth of untargeted mass spectrometry methods mirrors the growth in sequencing technologies and has generated extraordinary new insights into human and animal physiology^1,2^, disease processes^3,4^, microbial communities^5^, disease biomarkers^6^ and drug discovery^7^. Making sense of these large datasets, including spectral matching, statistical analysis, and machine learning, requires mathematical and statistical approaches to identify data patterns in noisy data to understand the biology. Missing values are quite common in large untargeted metabolomics datasets and can comprise up to 50% of the dataset and affect as many as 80% of the variables^8,9^. There are both biological and technical reasons why values may be absent. A metabolite in a sample may be missing due to: (1) an ion of a molecule being absent or below the limit of detection; (2) technical issues such as ion suppression^10,11^; (3) variability in sample processing ^12^; (4) variations in stability of molecules^13,14^; or (5) differences in ionization efficiencies^15^.

Missing values compromise the completeness of data which undermines the reliability of both univariate and multivariate statistical analyses such as fold-change analysis, *t*-tests, Analysis of Variance (ANOVA), regression-based analyses, and Principal Component Analysis (PCA)^16^. Additionally, missing values result in the loss of critical biochemical information, impeding the identification of expression patterns, biomarkers, the understanding of biological pathways, and their interactions^17,18^. Because many instances of missing values represent false negatives, dozens of methods for imputing missing values have been developed^19,20^. While variable in their approaches, imputation methods model missing data based on the non-missing values in the dataset. For instance, methods such as k-Nearest Neighbors (kNN) and Random Forest (RF) derive missing metabolite values using the values that exist for that same metabolite in other samples. This suggests that the amount of available data (or inversely the amount of missing data, i.e., “missingness”) may have a strong impact on the accuracy of imputation.

Another factor that could significantly impact the accuracy of imputations is the type of missing data which, if incorrectly assumed or modeled, could lead to inaccurate imputation values^21^. In metabolomics data, researchers have identified three primary classes of missing data: Missing Completely at Random (MCAR), Missing at Random (MAR), and Missing Not at Random (MNAR). Each mechanism has distinct characteristics and implications for data analysis and imputation methods^22,23^. Data points are MCAR when the probability of missingness is the same for all observations. MCAR data points are a random subset of the complete data, which means that analyses performed on the observed data can be unbiased. Simple imputation methods, such as minimum, mean or median imputation, can be appropriate in this case (^23–25^). In the context of metabolomics, data points would be MCAR in cases where some technical replicates detected signals, while others did not. Data is MAR when the probability of missingness depends only on the observed data and not on the missing data itself. An example of MAR data within metabolomics would be cases where undetected signals could be explained by the presence or absence of signals for another metabolite (e.g., inability to deconvolute co-eluting compounds)^26^. When data are MAR, more sophisticated imputation methods are needed, such as multiple imputation or maximum likelihood methods, that leverage the relationships within the observed data to account for the missingness^27^. Data is MNAR when the probability of missingness is related to external factors outside the features within the dataset. The most common example of MNAR data in metabolomics is the absence of signals at the limit of detection. Other MNAR could be due to differences related to sample type. MNAR imputation is the most challenging, as it requires assumptions about the unobserved data or external information to model the missingness mechanism^28^. In practice, it is very difficult to distinguish MCAR and MAR^29^, with some researchers advocating for alternative viewpoints for missing data altogether^30^.

An additional important aspect of metabolomics analysis is the issue of compositionality. Metabolomics, especially untargeted metabolomics, can have both compositional and non-compositional character. In many fields, including metabolomics, data may exist in a compositional form, meaning that the data represents parts of a whole. Compositional data are defined as constrained to sum to a fixed value (e.g., 1 or 100%), making the values within one file interdependent. In other words, if one value in compositional datasets increases all the other values must decrease, making the values non-independent (in technical terms, the data lie in a simplex rather than in Euclidean space). Alleviating this problem requires specialized transformations such as the centered log-ratio (CLR), additive log-ratio (ALR), or the isometric log-ratio (ILR), which convert the compositional data to real number space. Traditional statistical methods assume that data points are independent and can vary freely, but the inherent constraints of composition data create dependencies between the components, leading to potential pitfalls in data interpretation, including spurious correlations and misleading patterns^31^.

In untargeted metabolomics, the absolute value of the peak area can reflect the concentration of an analyte present in the sample. When not normalized to a fixed total, these values exhibit a non-compositional character because they are independent of the presence of other metabolites in the analysis. However, the distinction between compositionality and non-compositionality can sometimes be ambiguous, often depending on how the data is processed and interpreted. For instance, consider a fixed-volume plasma sample where 100 metabolites are detected. In a different sample, the same 100 metabolites are present along with a highly abundant drug. If one normalizes each sample to 100%, a common practice in untargeted metabolomics, all other metabolite signals in the second sample are proportionally reduced due to the presence of the drug. This normalization does not reflect a true decrease in their absolute quantities, thus introducing a compositional bias despite the original concentrations remaining unchanged. Another example of this complexity arises from ion suppression, where the introduction of certain molecules can hinder the detection of other metabolites. In this scenario, the measured abundance of some metabolites may be artificially lowered, complicating the interpretation of compositional versus non-compositional data. Further complicating this discussion are the structural vs observational zero values. Structural zeros indicate inherent limits in what is present, influencing the modeling approach and interpretation of results, while observational zeros reflect gaps in instrument sensitivity (peak detection settings) that can skew the perceived composition and relationships among components. Recognizing these impacts helps in the analysis and interpretation in compositional studies.

While raw metabolomic data can comprise a mix of compositional and non-compositional values, metabolite data becomes truly compositional after Total Ion Count (TIC) normalization, also known as relative abundance normalization. TIC normalization is commonly used with metabolomics data sets, particularly in mass-spectrometry based studies^32,33^. TIC normalization corrects for variations in overall signal intensity that may arise from differences in sample loading, ionization efficiency, or instrumental sensitivity during data acquisition^34^ These variations can introduce significant bias and obscure true biological differences between samples. TIC normalization mitigates these technical artifacts by scaling the signal intensities of individual metabolites relative to the total ion current^35^. Critically, TIC normalization adjusts the raw signal intensities of metabolites relative to the total ion current of the sample, effectively converting the data into a composition. TIC normalization ensures that the metabolite abundances are expressed as relative proportions of the total signal, making them comparable across different samples.

Understanding and applying compositional data analysis (CoDA) in metabolomics is critical because it allows for more accurate interpretation of the biological significance of relative metabolite abundances. By recognizing the compositional nature of TIC-normalized data, researchers can apply appropriate transformations, such as the CLR, to analyze the data without introducing biases or misinterpretations. The CLR transformation^36^ is a straightforward CoDA transformation that can be directly used in numerous statistical analyses and machine learning approaches, and is frequent applied in the fields of geology, molecular biology, and microbial ecology^37–39^. CLR transformation addresses the non-independence issue of compositional data by converting the data into real number space^39^. With the CLR transformation, each data point in a sample is replaced by the log-ratio of that sample value to the geometric mean, a form of averaging of the data from that sample, ensuring that all data points are centered around a mutual reference point. Another useful application of CLR transformation for metabolomics arises in its ability to facilitate comparative data analysis. Researchers are increasingly combining data from multiple ‘omics approaches (metagenomics, metabolomics, transcriptomics) to determine potential associations among them. Often, however, the distribution and scales of the values among these data sets can be remarkably different, and many statistical tests assume or require similar distributions across data. Using a common transformation ensures that all the datasets to be compared or integrated have a similar distribution and scale for comparisons.

In this study, we examined the effects of missing values on imputation using two of the most commonly used methods, namely k-Nearest Neighbors (kNN) and Random Forest (RF) using two unsimulated metabolomic datasets. The kNN and RF methods are frequently used in the literature and are often included in software packages. They have also been the most extensively validated in prior studies using simulated data^20^. While there are dozens of methods available for imputing data^40^, our goal was not to compare and contrast all the different imputation approaches. Rather, the focus of this paper was on the general impact of missing values and compositional data analysis on imputation. Specifically, our goals were to determine (1) how overall levels of missing data generally impact imputation and analytic interpretation of metabolomics data; and (2) how these imputation methods perform with very different types of missing data, namely MNAR and MCAR. Analysis with and without imputation were performed using raw data, TIC normalized data, and two types of CoDA transformation, CLR and the Robust CLR (RCLR) transformation^41^, and we examined the impacts of imputation at the whole sample-level and at the level of individual metabolites. Our results showed that both the amount and type of missing data had a profound impact on imputation and downstream interpretation, particularly at the level of individual metabolites. The impact also varied considerably depending on the type of missingness, with imputation performing especially poorly with MNAR data, and there was a strong correlation between the amount of missing data and the inaccuracy imputations. Handling missing data is a crucial step when working with metabolomics data and introduces additional complexity, as imputation methods may introduce bias and affect downstream interpretations^42^. Addressing missing values is paramount for maintaining the integrity and validity of untargeted metabolomics research, ensuring reproducible results, and facilitating meaningful biological insights. Overall, our findings caution against the blind use of imputation methods, especially with high levels of missingness and without a complete understanding of the types of missing data inherent in a dataset.

## RESULTS AND DISCUSSION

Several studies have already evaluated the accuracy of various imputation methods on untargeted metabolomics analysis. These studies largely rely on simulated datasets and with low numbers of missing values. However most untargeted metabolomics data have a lot of missing values and thus the results should be evaluated with few to large numbers of missing values. Here, we explore the effectiveness of commonly used imputation methods on two unsimulated datasets. In particular, we assessed how variation between samples changes, examined the effects of imputations when data is MCAR vs MNAR missing at the metabolite level, and evaluated the accuracy of imputations when missing values are introduced randomly and the true value is known. The first dataset was targeted, which included metabolites that were spiked into the samples at fixed concentrations along a logarithmic gradient. The second NIST dataset was untargeted and examined the metabolite profiles of homogenized human fecal samples which had either a vegetarian or omnivorous diet. The effect of the different imputations was assessed in conjunction with common normalization/transformation methods such as TIC normalization, CLR transformation, and RCLR transformation. Our key findings show that machine learning (ML) based imputations imputed missing values close to the detected signals within samples when the proportion of missing data across samples for a metabolite is low (e.g., 10% missing values). However, ML imputations cannot effectively handle MNAR data. Furthermore, we show that the proportion of missing values had a strong impact on the accuracy of imputation which affected the interpretation of the results.

### How do compositional transformations and imputations affect targeted metabolomics data where data is MNAR?

The first dataset we examined had data which was MNAR because it included metabolites spiked into an untargeted dataset at known concentrations (i.e., not random) that should be detected following the concentration gradient. However, some signals were not detected at the lowest concentrations for a subset of the metabolites due to limits of detection. Spiked samples are commonly used as internal standards and should not have missing signals, giving motivation to impute these missing values because we can guarantee the presence of the metabolite at a given concentration. The use of compositional transformations on the TIC normalized data is appropriate because all the individual ion counts are divided by the TIC, which sums to a fixed value (1.0 or 100%). While compositional analysis of the TIC-normalized data is clearly warranted, we also recognize that researchers also directly transform and impute using the raw data, often with similar results. Therefore, we have repeated the CoDA below with the raw data and included these results as supplemental for comparison.

As mentioned in the introduction, several mathematical transformations have been developed to resolve this compositionality issue including the CLR and more recently the RCLR, both of which convert compositional data into real number space alleviating the non-independence issue. Since both the CLR and RCLR are log-ratio transformations, and the log of zero is undefined, zero handling becomes an important issue. For the CLR, zeros are replaced prior to transformation with a very small number, while the RCLR removes the zeros during the transformation and then replaces them with zeros which are often converted to a small number after transformation for downstream analysis.

To assess the impacts of normalization and compositional transformation on MNAR data we compared the PCA clustering of samples at increasing concentrations without imputation.

**Figure 1 A-D** shows PCA analyses for the targeted dataset with no transformations (raw data), TIC normalization, TIC-CLR transformation (TIC normalization followed by CLR), and TIC-RCLR transformation (TIC normalization followed by RCLR) without machine learning imputations (0 values substituted with minimum value following transformation). The total variation explained between the different methods was similar. PCA for TIC-CLR transformed data explained the most variation with the first two principal components (55.05%; **Figure 1B**), while the PCA for the raw data captured the least (53.78%; **Figure 1A**). PCA for TIC-CLR transformed data captured more variation on the second principal component compared to other methods (**Figure 1C**). Clustering of replicates within conditions was similar across the different methods, with the exception of the TIC-CLR transformed data, which exhibited different clustering compared to the other three approaches (**Figure 1C**). It is worth noting that selecting varying values of pseudocounts for the TIC-CLR transformation produces different clustering of samples.

**Figure 1.**
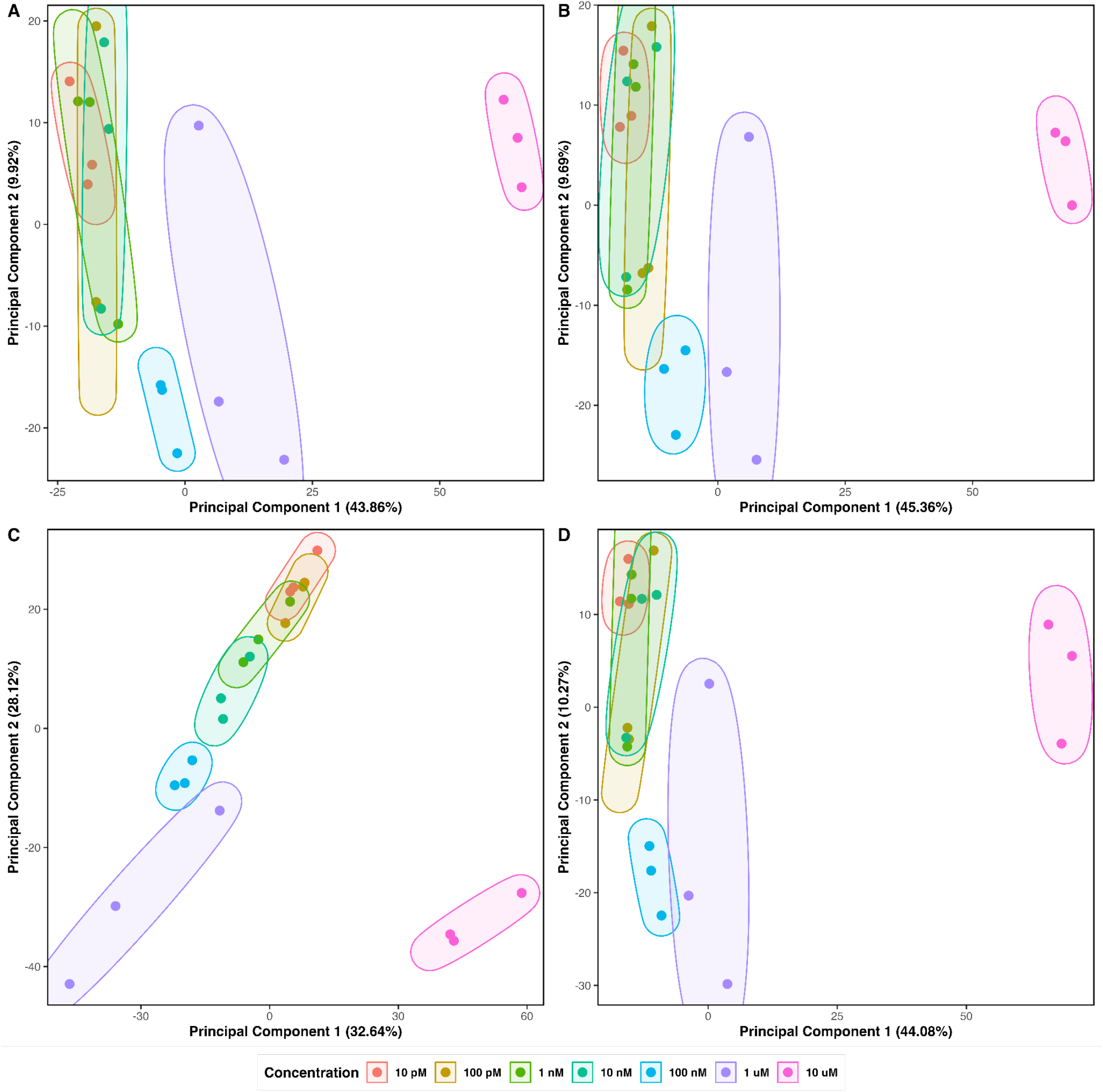
PCA plots for different transformations on targeted metabolomics data; **A**. PCA plot for raw data, **B**. PCA plot for TIC normalized data, **C**. PCA plot for TIC-CLR transformed data (pseudocount = 1e-12), **D**. PCA plot for TIC-RCLR transformed data, followed by minimum value imputation.

However, since the TIC normalized data contains many proportions with very small values, a pseudocount smaller than all TIC proportions was appropriate (1e-12). The same analysis performed after transforming the raw data instead of the TIC-normalized data produced similar results for the RCLR-transformed data, although the CLR explained more of the variation in the first two principal components (**Supplemental Figure 1**).

To determine the effects of imputation methods on PCA clustering and visualization, we then applied two commonly used imputation methods, kNN and RF. We compared cases where TIC-normalized data was first imputed, then CLR transformed, and RCLR-transforming the TIC-normalized data, then imputing missing values for the unaffected 0’s. Since imputation can be performed either before or after transformation, we examined both possibilities. **Figure 2A-D** shows PCA analyses for the targeted dataset using machine learning imputations for missing data with TIC-RCLR and TIC-CLR transformations. Since CLR transformation does not accept missing values (one cannot compute log(0)), missing value imputation was performed on TIC normalized data prior to CLR transformation. For the TIC-RCLR transformation, imputation was performed after the transformation because the missing/0 values are ignored during the process. Similar to the results in **Figure 1**, there were minimal differences in the total variation explained between the different approaches. PCA for TIC-RCLR transformation followed by kNN imputation explained the most variation with the first two principal components (42.23%; **Figure 2B**), while RF imputation on TIC normalized data followed by CLR transformation captured the least (38.88%; **Figure 2C**). Clustering of replicates within conditions was also highly similar; only RF imputation on TIC normalized data followed by CLR transformation exhibited different clustering, showing reduced resolution of groups (**Figure 2C**). While the results of kNN and RF imputations were similar, the variation explained using the imputed data was consistently lower than the non-imputed analyses. Similar results were found with transformations performed on the raw data (**Supplemental Figure 2**).

**Figure 2.**
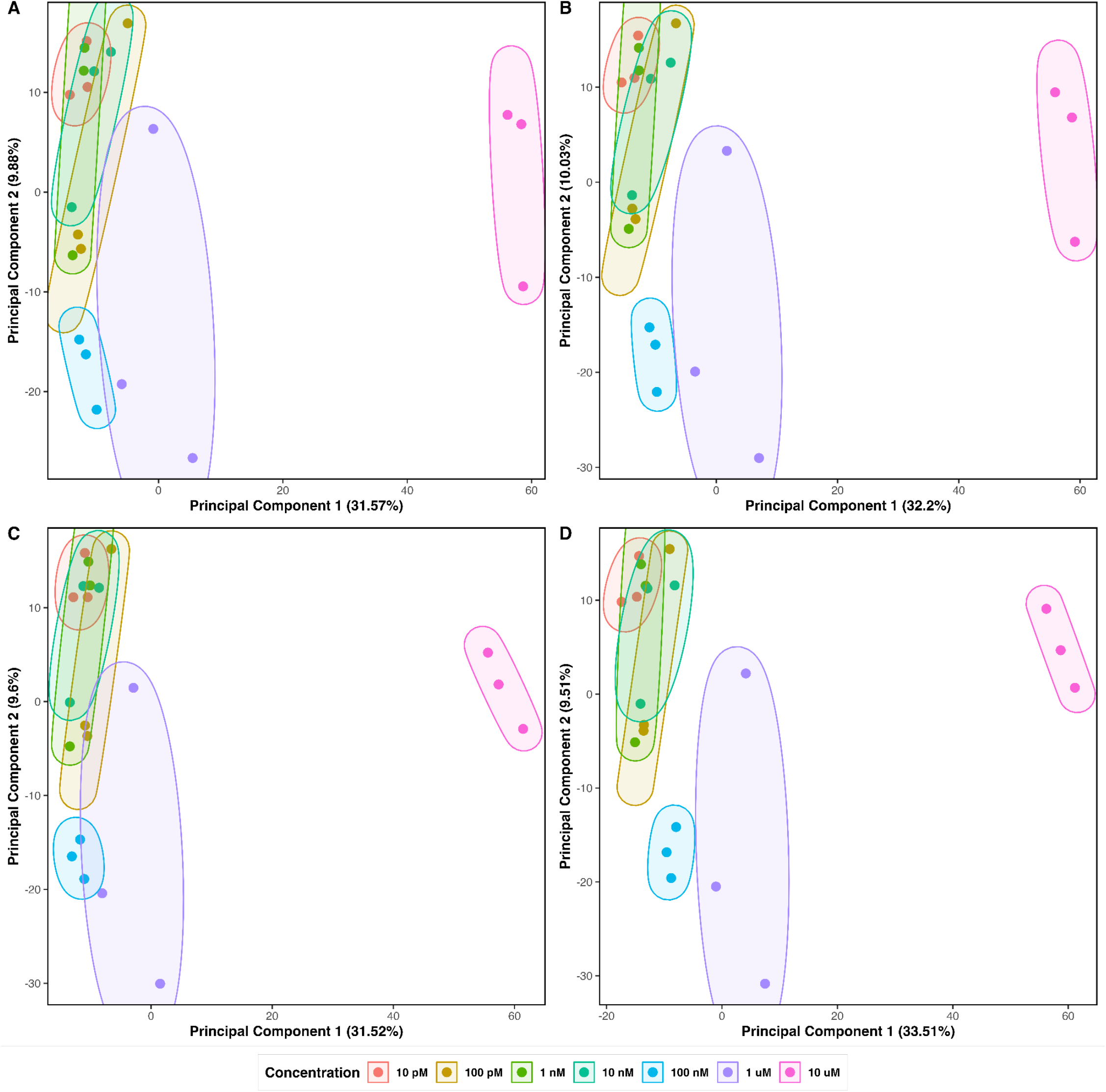
PCA plots of the targeted metabolomics data using machine learning imputations and two different log-ratio transformation methods. **A**. TIC-RCLR transformed data, followed by RF imputation, **B**. TIC-RCLR transformed data, followed by kNN imputation, **C**. RF imputation on TIC normalized data, followed by CLR transformation, **D**. kNN imputation on TIC normalized data, followed by CLR transformation.

To determine the effects of imputation methods on individual MNAR metabolite abundances, we determined the effects of kNN and RF missing-value imputations to non-imputed values on individual metabolites with different levels of missingness. **Figure 3A-F** shows boxplots for a metabolite-level comparison of transformations and imputations where data points are MNAR-type missing. All data points were missing for these spiked-in standards at the two lowest concentrations. TIC-CLR transformation preserved the log-scale proportional relationship between data points in TIC-normalized data because the data is converted into log-ratios relative to the geometric mean (**Figure 3B** and **Figure 3C**). The CLR transformation was more similar to the TIC-normalized data, compared with the RCLR transformation with minimum value imputation (**Figure 3D**), though both preserved the relationships between the different concentration standards. Following TIC-RCLR transformation, both kNN and RF imputations failed to effectively impute missing values below the limit of detection (**Figure 3E** and **Figure 3F**). Both kNN and RF overestimated the missing data, with a larger overestimation using RF compared to kNN imputation (**Figure 3F**). Nearly identical results were determined when transformations were performed using raw data (**Supplemental Figure 3**).

**Figure 3.**
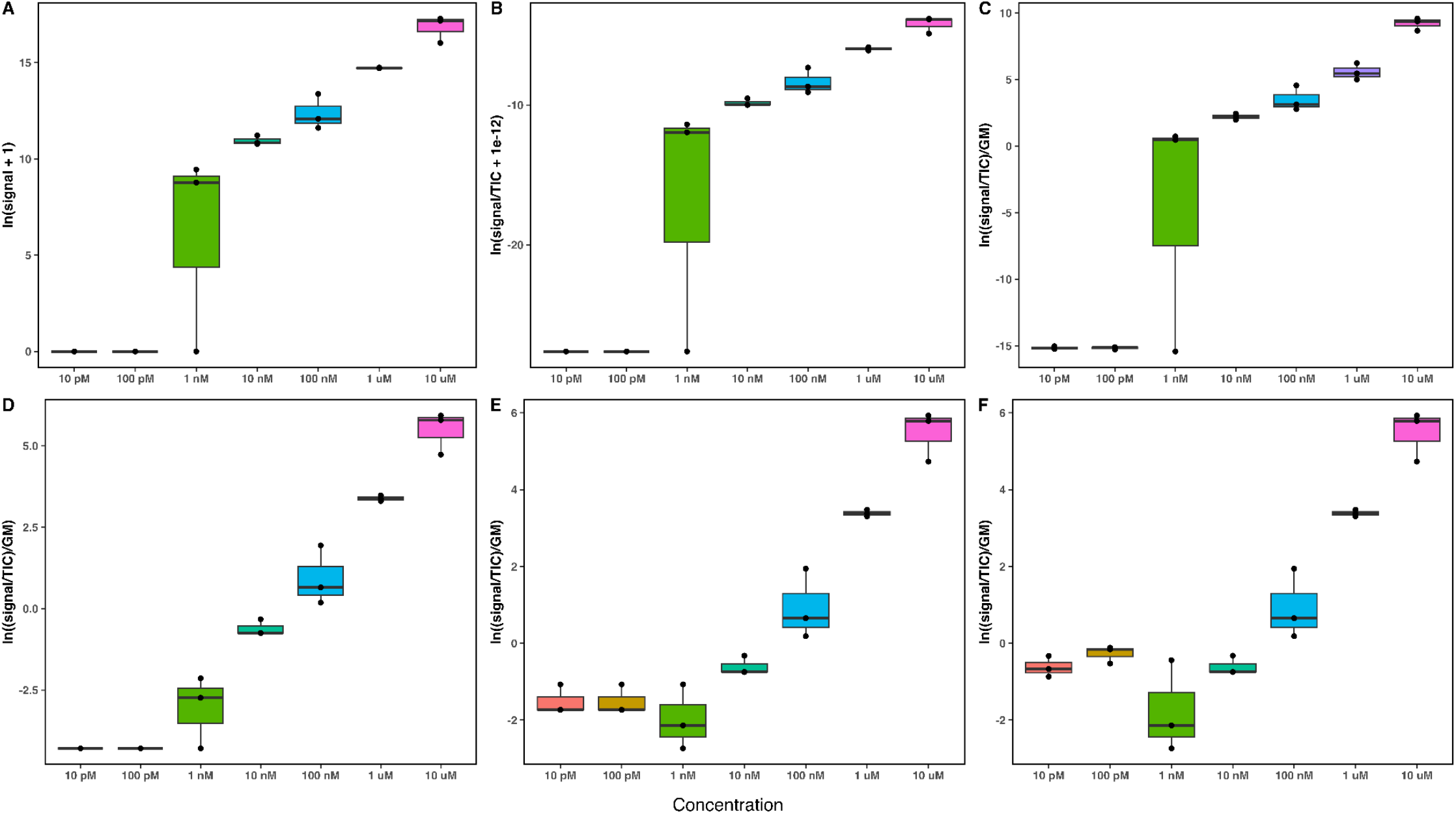
Boxplots of transformations and imputations for aspartame, where the 2 lowest concentration thresholds have all missing data; **A**. Raw data plotted on a natural log scale, **B**. TIC normalized data plotted on a natural log scale, **C**. TIC-CLR transformed data (pseudocount = 1e-12), **D**. TIC-RCLR transformed data, followed by minimum value imputation, **E**. TIC-RCLR transformed data, followed by kNN imputation, **F**. TIC-RCLR transformed data, followed by RF imputation.

**Figure 4A-F** shows another metabolite-level comparison with MNAR-type missing data. In this example, an additional two orders of magnitude are missing. Similar to **Figure 3**, TIC-RCLR transformation followed by kNN and RF imputations failed to effectively impute missing values below the limit of detection (**Figure 4E** and **Figure 4F**). RF overestimated missing values more than kNN imputation (**Figure 4E** and **Figure 4F**). Nearly identical results were determined when transformations were performed using raw data (**Supplemental Figure 4**).

**Figure 4.**
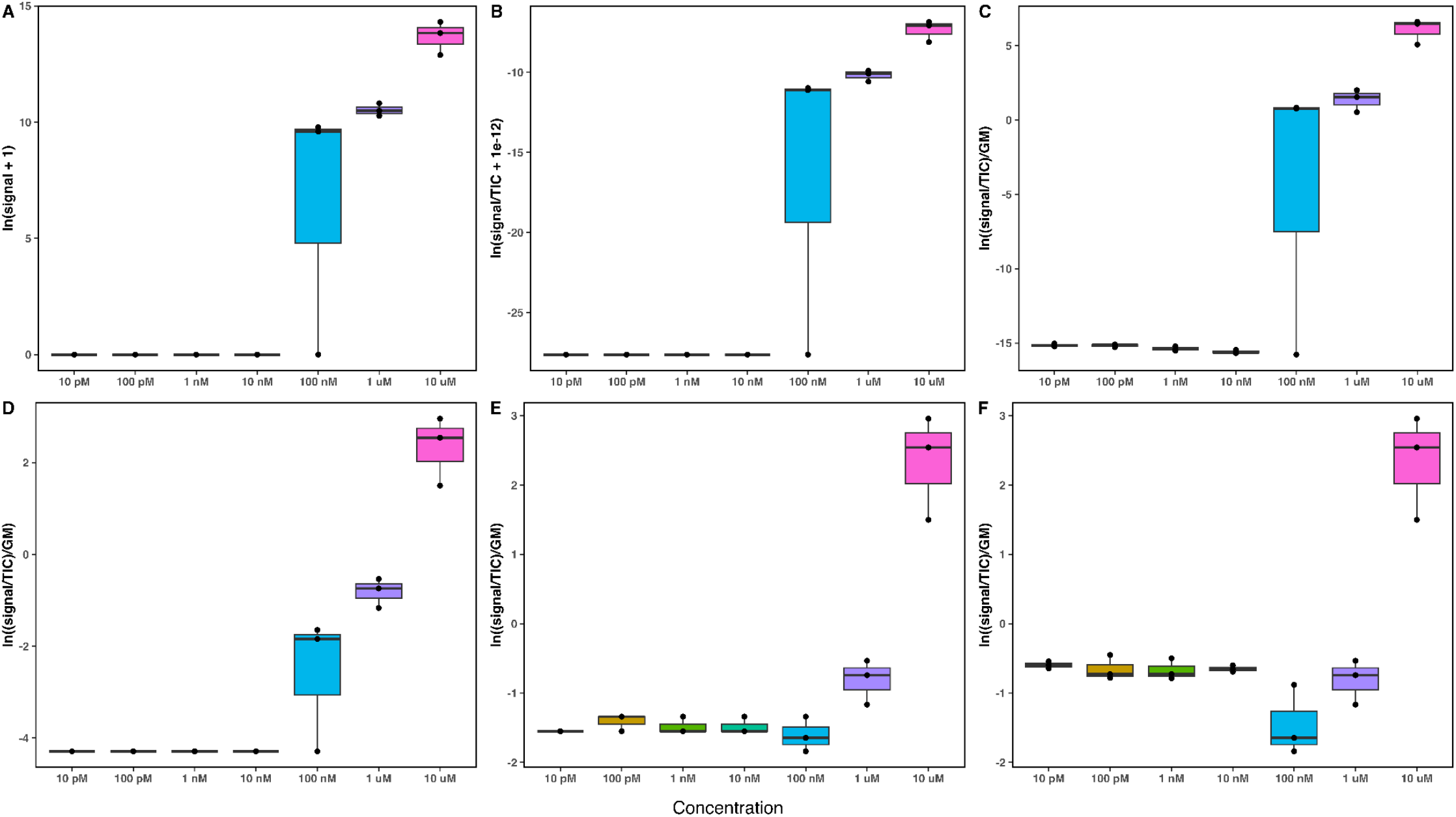
Boxplots of transformations and imputations for estrone, which is missing signals for the four lowest concentrations; **A**. Raw data plotted on a natural log scale, **B**. TIC normalized data plotted on a natural log scale, **C**. TIC-CLR transformed data (pseudocount = 1e-12), **D**. TIC-RCLR transformed data, followed by minimum value imputation, **E**. TIC-RCLR transformed data, followed by kNN imputation, **F**. TIC-RCLR transformed data, followed by RF imputation

**Figure 5A-D** shows model-based (logistic regression) imputation to handle MNAR-type missing data points. On the log scale, metabolites which had detected signals for all concentrations exhibited a linear relationship (**Figure 5B**). To impute MNAR missing values, a logistic regression model was fitted using only existing data points, then used to impute missing values (**Figure 5C**). Following logistic regression model-based imputation, imputed values conform to the expected pattern for data points with order of magnitude concentration differences (**Figure 5D**).

**Figure 5.**
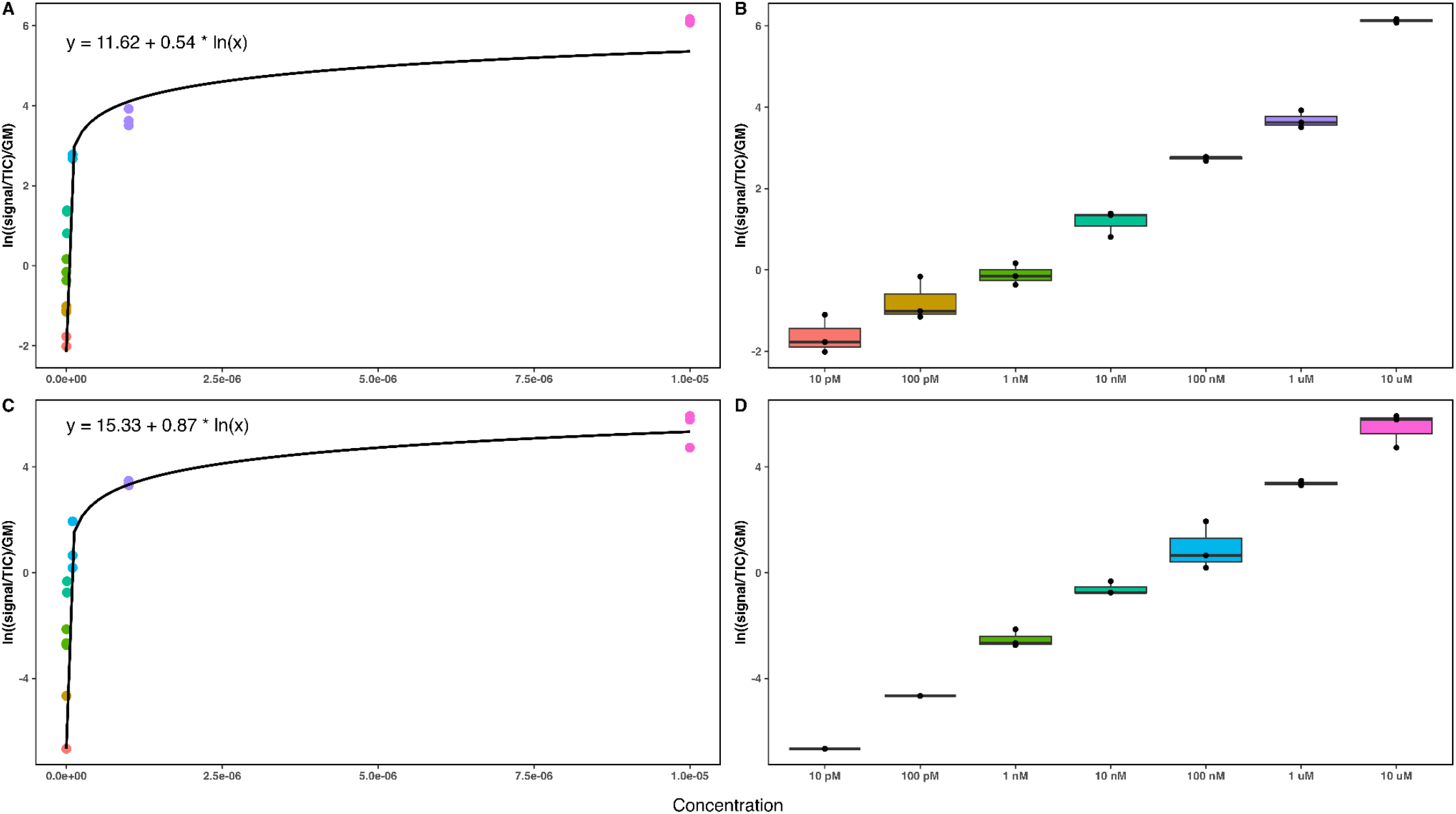
Regression-based modeling approach for handling missing values in targeted metabolomics data which conform to MNAR-type pattern; **A**. Logarithmic regression for a C17 sphingosine, which had detected signals at all concentrations and for all replicates (TIC-RCLR-transformed data), **B**. Boxplots for C17 sphingosine, **C**. Logarithmic regression and imputation for aspartame, which had missing signals at 2 lowest concentrations (10 pM and 100 pM) for all replicates (TIC-RCLR transformed data), **D**. Boxplot assessing logarithmic regression imputation following TIC-RCLR transformation for aspartame.

### How do compositional transformations and imputations affect untargeted metabolomics data which is both MCAR and MNAR?

The second dataset, a metabolomics analysis of people on a vegan or omnivore diet^43^, we examined was derived from samples and had missing values that were both MCAR and MNAR. The MCAR missing values had no discernable pattern to their missingness, which might have been a result of issues with sample handling or detection. However, we also found missing values that appeared to be MNAR because they were present in all the omnivore diet samples but absent in all the vegetarian diet samples (and vice versa). We expect many environmental datasets will have some combination of MCAR and MNAR values, and our results show that the type of missing data has a profound impact on imputation accuracy.

To assess the impacts of normalization and compositional transformation on MCAR/MNAR data, we compared PCA clustering of samples within the different diets without machine learning imputations (**Figure 6A-D**). PCA for raw data explained the most variation in the first two principal components (79.92%; **Figure 6A**), while PCA for the TIC-CLR transformed data captured the least (68.99%; **Figure 6C**). Clustering of replicates within conditions was similar; four discrete clusters were observed with raw data, TIC normalized data, and TIC-RCLR transformed data with minimum value imputation (**Figures 6A-B, 6D**). PCA for TIC-CLR transformed data failed to resolve clustering of samples with the same resolution (**Figure 6C**). Nearly identical results were determined when transformations were performed using raw data (**Supplemental Figure 5**).

**Figure 6.**
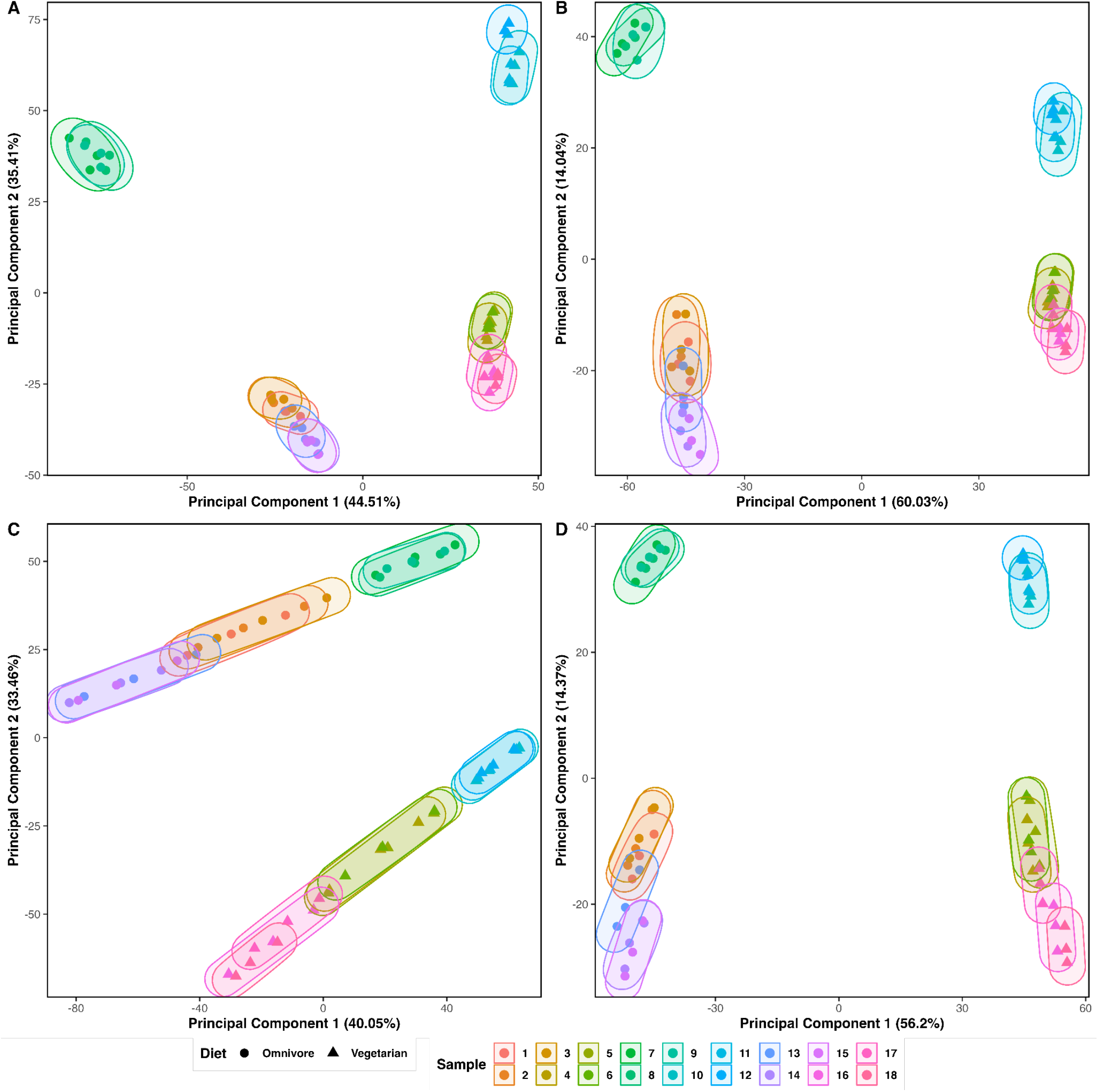
PCA plots for normalizations on untargeted metabolomics data with no machine learning based imputations; **A**. PCA plot for raw data, **B**. PCA plot for TIC-normalized data, **C**. PCA plot for TIC-CLR transformed data, **D**. PCA plot for TIC-RCLR transformed data, followed by minimum value imputation for missing values.

We then determine the effects of kNN and RF imputation methods on PCA clustering and visualization of MCAR/MNAR data. **Figure 7A-D** shows PCA analyses for the untargeted dataset using machine learning imputations on missing data with TIC-RCLR and TIC-CLR transformations. The total variation captured among the different methods was very similar.

**Figure 7.**
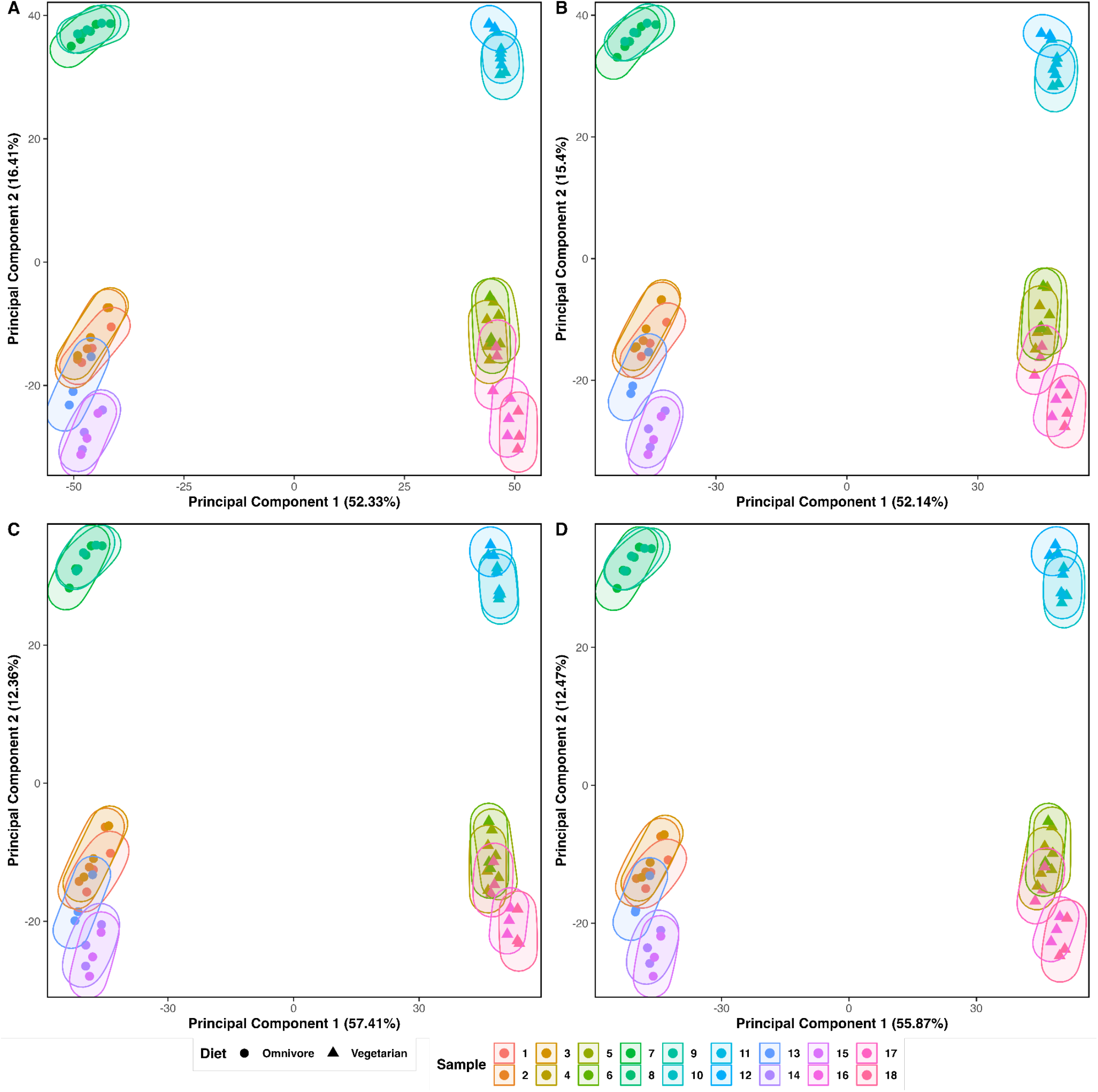
PCA plots of the untargeted metabolomics data using machine learning imputations and two different log-ratio transformation methods. **A**. TIC-RCLR transformed data followed by RF imputation, **B**. TIC-RCLR transformed data followed by kNN imputation, **C**. RF imputation on TIC normalized data followed by CLR transformation, **D**. kNN imputation on TIC normalized data followed by CLR transformation.

PCA for TIC-RCLR transformed data followed by kNN imputation captured the most variation on the first two principal components (68.74%; **Figure 7B**), while RF imputation on TIC-normalized data followed by CLR transformation captured the least (64.04%; **Figure 7C**). Clustering of replicates within conditions was highly similar in all four analyses. While the difference is smaller, we note that the variance explained using the imputed data was still consistently lower than the variance explained using the non-imputed data. Similar results were determined when transformations were performed using raw data (**Supplemental Figure 6**).

To test the hypothesis that imputation accuracy increases as the proportion of missing data across features decreases, we binned percentage of missing data within a feature into 3 discrete categories: features that had less than 10% missing values, features that had 50% missing values, and features that had greater than 90% missing values. For each threshold of missing data, we substituted another missing value for an entry that has a known value and assessed whether the accuracy of the ML-based imputation methods improves. **Figure 8A-D** shows distance comparisons between true value and imputed value for artificially produced missing values at different thresholds of missing data within a feature for untargeted metabolomics data. Whether missing data is imputed prior to transformation or after, the absolute distance between true value and ML-imputed imputed value varies inversely as the percentage of detected signals increases across samples for a metabolite.

**Figure 8.**
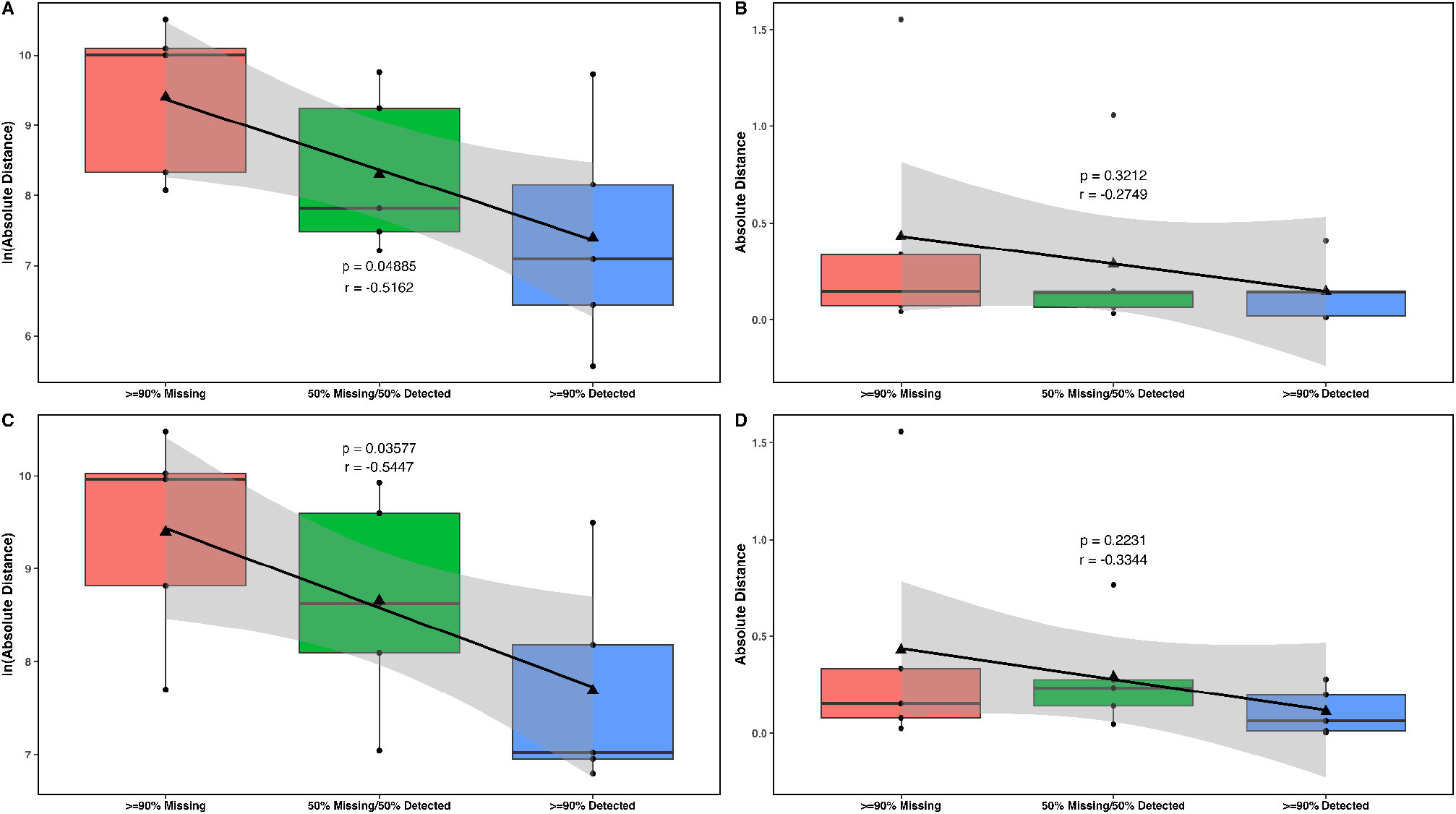
Distance comparisons between true value and imputed value applied to non-transformed untargeted metabolomics data and RCLR transformed data. **A**. kNN imputation for missing values on raw data, **B**. RCLR transformation on data, followed by kNN imputation for missing values, **C**. RF imputation for missing values on raw data, **D**. RCLR transformation on data, followed by RF imputation for missing values.

Finally, to determine the effects of imputation methods on the untargeted MCAR/MNAR metabolite abundances, we performed kNN and RF missing-value imputations on individual metabolites with different proportions of missing data. **Figure 9A-F** shows a metabolite-level comparison of transformations and imputations where data points are MCAR-type missing within the untargeted dataset using a metabolite with >90% detected signal across all samples. TIC-CLR transformation preserved the proportional relationship between data points observed in TIC-normalized data (**Figure 9B** and **Figure 9C**). TIC-RCLR transformation followed by kNN and RF imputations yielded highly similar results; both methods imputed missing values close to the detected signals (**Figure 9E** and **Figure 9F**). Nearly identical results were determined when transformations were performed using raw data (**Supplemental Figure 7**).

**Figure 9.**
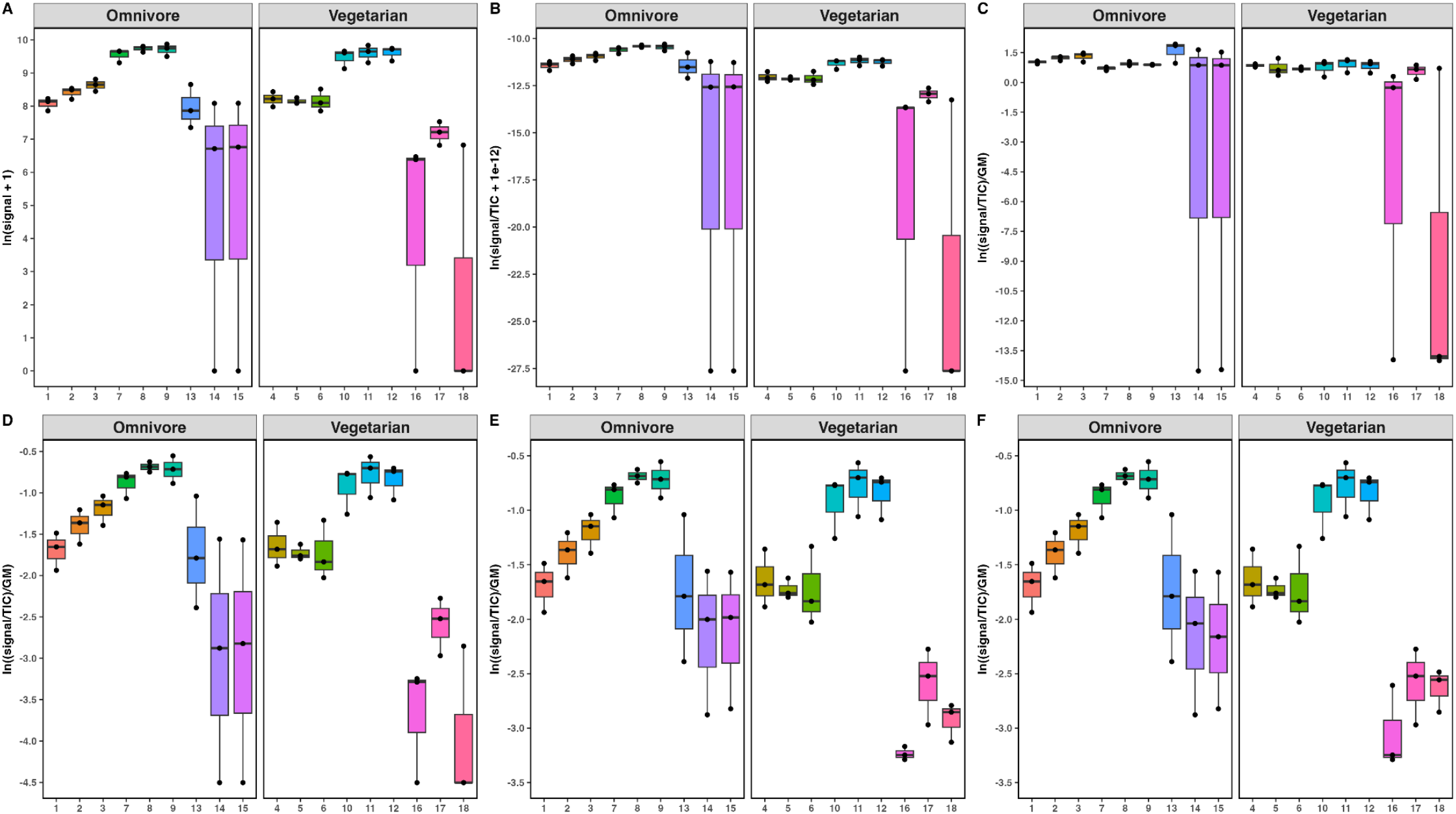
Boxplots of transformations and imputations for an unknown metabolite with >90% detected signals across all samples in the untargeted dataset; **A**. Raw data plotted on a natural log scale, **B**. TIC normalized data plotted on a natural log scale, **C**. TIC-CLR transformed data (pseudocount = 1e-12), **D**. TIC-RCLR transformed data, followed by minimum value imputation, **E**. TIC-RCLR transformed data, followed by kNN imputation, **F**. TIC-RCLR transformed data, followed by RF imputation.

**Figure 10A-F** shows a metabolite-level comparison of transformations and imputations where data points are MCAR-type missing within the untargeted dataset; the metabolite had 50% detected and 50% undetected signals across all samples, with missing values in both diet groups. TIC-RCLR transformation followed by minimum value imputation preserved the proportional relationship between data points observed in the TIC normalized data (**Figure 10B** and **Figure 10D**), while TIC-CLR transformation did not (**Figure 10C**) due to centering data points around the sample-specific geometric mean and bias introduced by 0-substitution with pseudocounts during transformation. TIC-RCLR transformation followed by kNN and RF imputation gave similar results; both methods imputed missing values close to the detected signals (**Figure 10E** and **Figure 10F**). Nearly identical results were determined when transformations were performed using raw data (**Supplemental Figure 8**).

**Figure 10.**
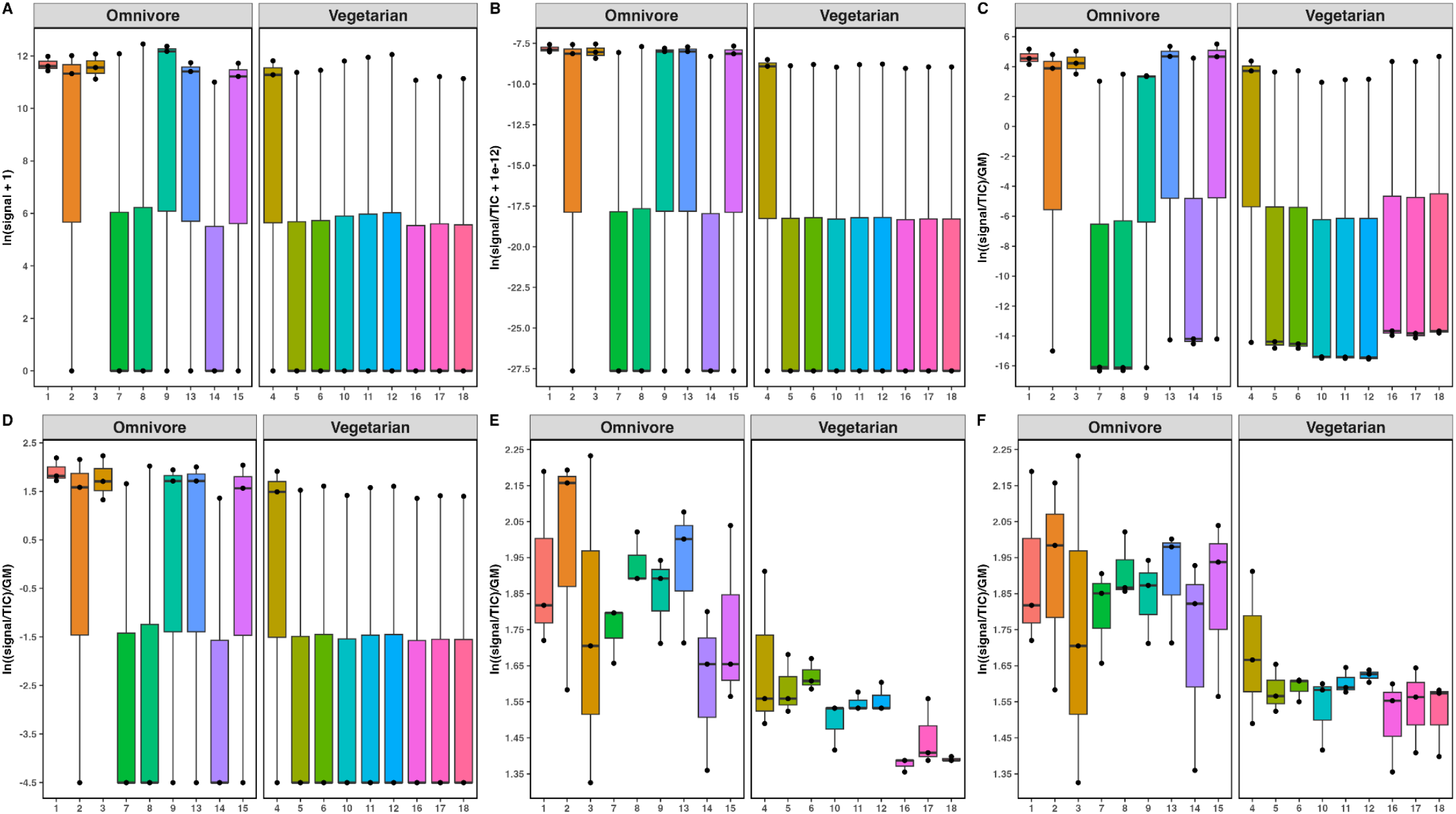
Boxplots of transformations and imputations for a metabolite with 50% detected signals, 50% missing signals across all samples in the untargeted dataset. **A**. Raw data plotted on a natural log scale, **B**. TIC normalized data plotted on a natural log scale, **C**. TIC-CLR transformed data (pseudocount = 1e-12), **D**. TIC-RCLR transformed data, followed by minimum value imputation, **E**. TIC-RCLR transformed data, followed by kNN imputation, **F**. TIC-RCLR transformed data, followed by RF imputation.

**Figure 11A-F** shows a metabolite-level comparison of transformations and imputations where data points are MNAR-type missing within the untargeted dataset. While the previous metabolites were MCAR, this example showcases MNAR data in the untargeted context because the metabolite had 50% detected and 50% undetected signals across all samples; however, undetected signals were only in the vegetarian diet samples. Due to the inherent nature of the CLR transformation, it is inevitable to introduce some bias with the transformation attributed to the pseudocount; even with a very small pseudocount (1e-12), interpretations could lead to misleading results on account of dividing by the geometric mean of each sample during the transformation (**Figure 11C**). TIC-RCLR transformation followed by minimum value imputation preserved the proportional relationship between data points observed in the TIC-normalized data (**Figure 11B** and **Figure 11D**) while TIC-CLR transformation did not (**Figure 11C**). Both kNN and RF imputations produced completely fabricated results on account of the fact that there was no data to impute from within the same group of individuals (vegetarian) – the only detected signals for the metabolite all came from samples which had an omnivore diet (**Figure 11E** and **Figure 11F**). Naively, the imputations give the illusion that the metabolite profile was similar amongst individuals in both diets. However, the metabolite was completely undetectable with individuals on a vegetarian diet. Nearly identical results were determined when transformations were performed using raw data (**Supplemental Figure 9**).

**Figure 11.**
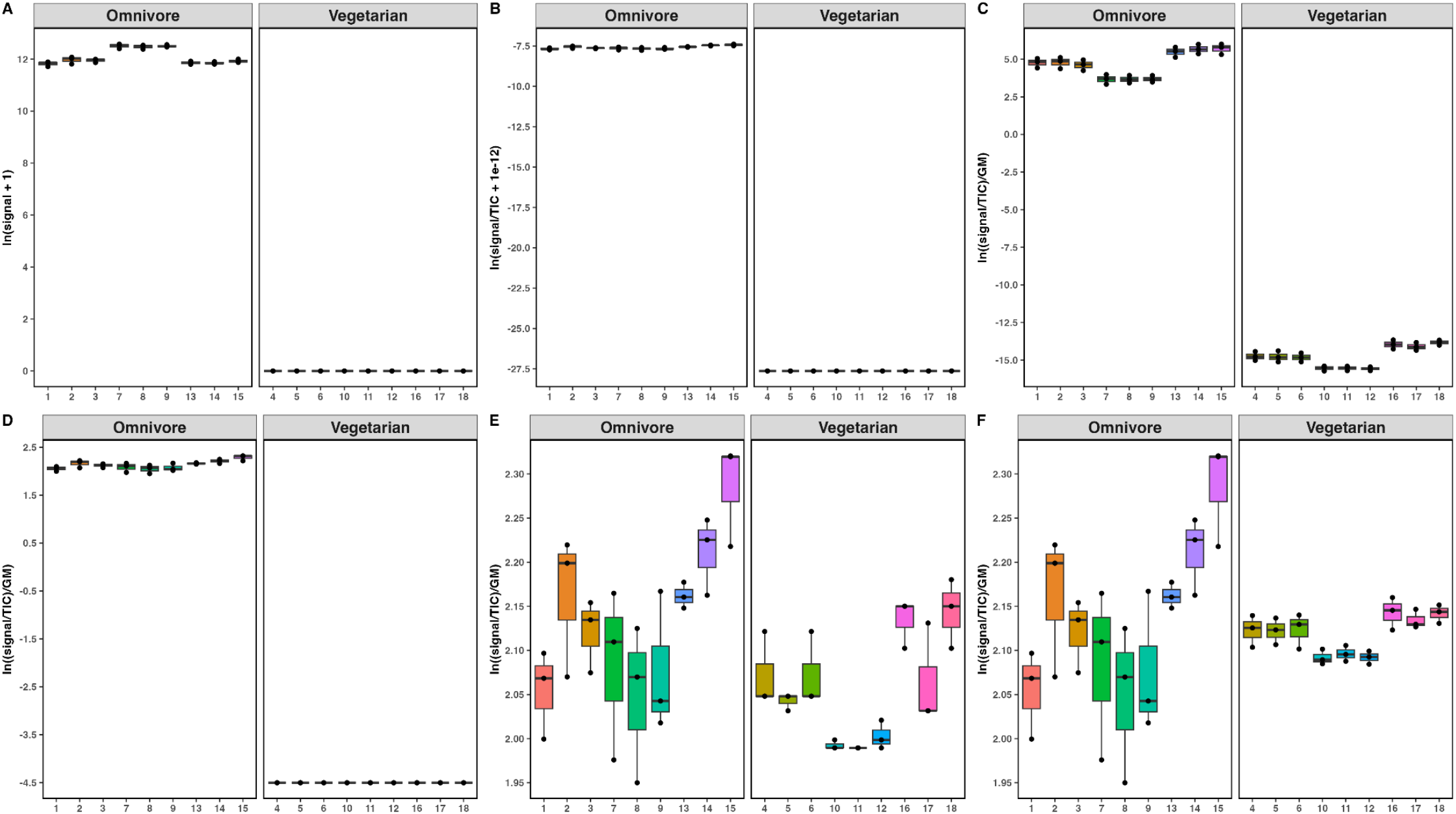
Boxplots of transformations and imputations for a metabolite with 50% detected signals, 50% missing signals across all samples in the untargeted dataset. **A**. Raw data plotted on a natural log scale, **B**. TIC normalized data plotted on a natural log scale, **C**. TIC-CLR transformed data (pseudocount = 1e-12), **D**. TIC-RCLR transformed data, followed by minimum value imputation, **E**. TIC-RCLR transformed data, followed by kNN imputation, **F**. TIC-RCLR transformed data, followed by RF imputation.

**Figure 12A-F** shows a metabolite-level comparison of transformations and imputations where the metabolite had <10% detected signals across all samples. TIC-RCLR transformation followed by minimum value imputation preserved the proportional relationship between data points observed in the TIC-normalized data (**Figure 12B** and **Figure 12D**) while TIC-CLR transformation did not (**Figure 12C**). TIC-RCLR transformation followed by kNN and RF imputation gave different results; RF imputation consistently had greater imputed values when compared with kNN imputation, a similar finding when compared with the targeted dataset (**Figure 12E** and **Figure 12F**). Both RF and kNN imputed missing values gave the illusion that there were discernible differences between diets and across diets; however, this metabolite was almost completely undetectable across all samples. Nearly identical results were determined when transformations were performed using raw data (**Supplemental Figure 10**).

**Figure 12.**
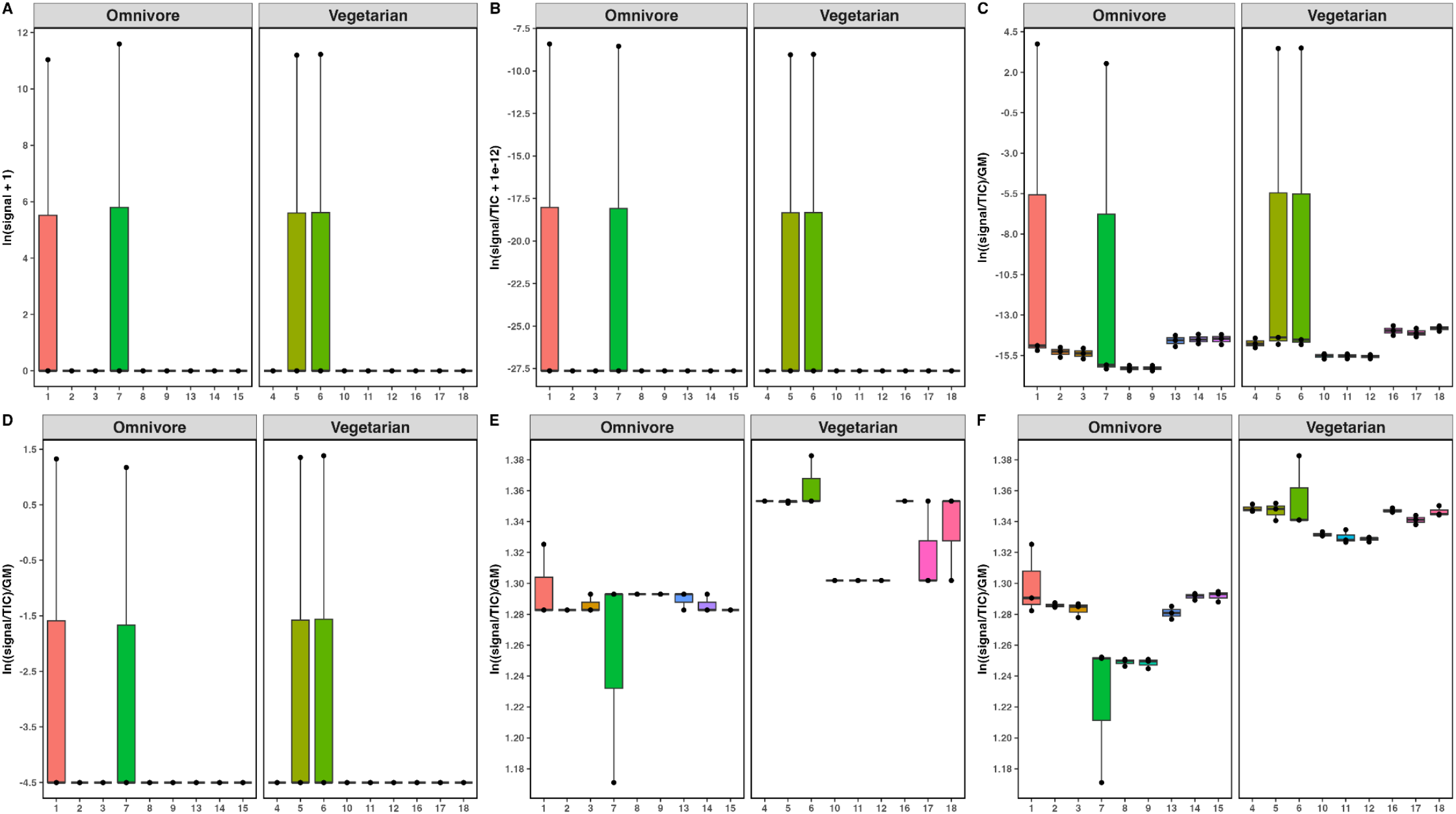
Boxplots of Transformations and imputations for a metabolite <10% detected signals across all samples in the untargeted dataset. **A**. Raw data plotted on a natural log scale, **B**. TIC normalized data plotted on a natural log scale, **C**. TIC-CLR transformed data (pseudocount = 1e-12), **D**. TIC-RCLR transformed data followed by minimum value imputation, **E**. TIC-RCLR transformed data followed by kNN imputation, **F**. TIC-RCLR transformed data followed by RF imputation.

Given the clear impact of missing data on imputation accuracy in our unsimulated targeted and untargeted datasets, we directly explored the impacts of different levels of missingness on MCAR imputation accuracy. **Table 1** summarizes a comparison of RF and kNN based imputations where data is MCAR. RF imputation consistently outperformed kNN imputation in terms of accuracy assessed by RMSE. In both cases, the accuracy declines as the percentage of missing data increases.

**Table 1.** Normalized Root Mean Square Error assessment of RF and kNN imputations at different thresholds of randomly assigned missing data for untargeted metabolomics.

| % Missing | NRMSE: TIC-RCLR followed by RF | NRMSE: TIC-RCLR followed by kNN |
| --- | --- | --- |
| 20% | 0.3405 | 0.3701 |
| 40% | 0.3503 | 0.3864 |
| 60% | 0.3678 | 0.4071 |
| 80% | 0.4136 | 0.5284 |

## Conclusion

In this study, we analyzed the effects of imputation on metabolomics data using several widely recognized methods: simple substitution, k-Nearest Neighbors, and Random Forest. The primary objective was to understand how these imputation techniques were affected by levels of missing data and by the types of missing data, e.g., MCAR and MNAR. Our findings showed that both kNN and RF imputation methods worked well at low proportions of missing data across samples and when the data was MCAR. We also showed that compositional data methods performed well across datasets, with little evidence of distortion compared to raw and TIC-normalized methods. However, the accuracy of kNN and RF imputations declined markedly as the proportion of missing data increased in both MCAR and MNAR data, and both performed especially poorly with MNAR data. At very high levels of missing data, and when data is MNAR, kNN and RF imputations falsely imputed differences in metabolite abundance between samples that did not exist in the real data. The effect of overestimation was particularly dramatic with individual metabolites, but the effect was also observed in the PCA analyses of both targeted and untargeted datasets where imputation resulted in a loss of explanatory power. With all methods, it is important to note that imputation often overestimated the abundance of a molecule’s signal if it is truly absent from the data even in the case of minimum value imputation.

Our finding that higher levels of missingness in MCAR data corresponded to poorer imputation accuracy makes sense considering that these ML methods train on the existing values in the data set. The more data available for training, the more accurate the imputation. In some cases, we discovered that the methods were imputing large numbers of values after training on only a single existing data point. While we only tested the effect of missing data with two of the many dozens of available imputation methods, similar problems likely plague any imputation approach: the fewer the values available for training or estimation, the lower the imputation accuracy. Our discovery that imputation methods performed considerably worse with MNAR also makes sense when we consider that the underlying assumptions of these methods is that the missing data have the same distribution as the non-missing data. Indeed, the imputation methods were highly accurate when we accurately modeled the MNAR data for the targeted dataset with a logistic regression model. Identifying the pattern of MNAR in the untargeted data set also showed how imputation assumptions were clearly violated as they assumed the pattern of missingness was the same for both omnivore and vegetarian samples.

While imputation techniques offer a seemingly convenient solution for handling missing data, they present several significant drawbacks that warrant cautious consideration. The variability inherent in different imputation approaches introduces a degree of uncertainty that can undermine the reliability of the research findings. Imputations are fundamentally based on the existing data. The less data available for imputing, the worse the imputations. As a result, the imputed values might not accurately reflect the true underlying patterns, thereby compromising the integrity of the analysis. This is particularly problematic in cases where the missing data mechanism is not fully understood or is incorrectly assumed to be missing at random (MCAR or MAR) when it is not (MNAR), or vice versa. Our results with real datasets show that the erroneous assumption of the missing data mechanism may lead to inappropriate imputation methods, further exacerbating inaccuracies in the dataset. If one is sure of the classification of missingness, such as with MNAR with log-scale spiked in standards, then an appropriate imputation method can be applied with confidence. However, in many untargeted datasets, the type of missingness may be unclear or even mixed, suggesting great caution should be used with the blind use of imputation methods.

Imputation may work well for data where most of the signals are shared. Such experimental designs might include analysis of organisms where a single gene is deleted, or a single new member is added to a panel of bacteria, or datasets such as plasma metabolites where the same metabolites are observed very frequently (e.g. 50-100%) across the samples. These study designs tend to reflect central core metabolism of the samples that are being investigated. However, as factors like diet, microbiome variability, environmental exposures, and medication use differ among individuals, many metabolites may be present infrequently—often in less than 10% of samples within a cohort. Thus, one may have to make choices about whether to perform data imputation. If one filters the data for only common features, as is often done in statistics for untargeted metabolomics data, it is imperative to be transparent and report which data were included and which were excluded. This transparency helps ensure that the analysis remains interpretable and that potential biases are clearly communicated.

Reproducibility, a cornerstone of scientific research, is also threatened by the use of imputations. Imputation methods often rely on random processes, model-based predictions, or machine learning techniques. Different researchers might produce different imputed datasets from the same original data, leading to variability in results. This lack of reproducibility undermines the credibility of the research and makes it difficult for other scientists to validate or build upon the findings. A prime example of this is demonstrated with machine learning approaches, which typically rely on a pseudorandom number to seed the results (i.e., set.seed() in R and random_state in Python) and be reproducible. Changing these numbers could lead to vastly different results particularly in the case of Random Forest imputation and other tree-based methods.

Ultimately, imputations involve fabricating data that does not exist, which raises ethical and methodological concerns. By introducing artificial data into the analysis, researchers risk presenting findings that are not truly reflective of the real-world phenomena they aim to study. Therefore, it is advisable to consider alternative strategies for handling missing data, such as robust statistical methods that can accommodate missing data without the need for imputation, where possible designing studies that minimize the occurrence of missing data or have increased number of biological/technical replicates, and compositional data transformations. In light of these considerations, it is prudent to approach the use of imputations with great caution^44^. Ensuring the validity, reliability, and reproducibility of research should take precedence, guiding researchers towards more transparent and scientifically sound methods for managing missing data.

## METHODS

### Tandem Mass Spectrometry Analysis

We used two publicly-available untargeted metabolomics datasets in our analyses. The details of data collection for each dataset are presented below.

#### Targeted dataset

Dataset 1 was collected as described in Melnik et al.^45^. The dataset contains a mixture of 41 standards spiked in at equimolar concentrations to 10 uM fecal extract. Standards were added across a concentration gradient of 10 pM, 100pM, 1 nM, 10 nM, 100 nM, 1 uM, and 10 uM with three replicates per concentration. Briefly, MS/MS data was acquired in positive mode on a Q Exactive Orbitrap (Thermo Fisher Scientific, Waltham, MA) using data-dependent acquisition. Samples were separated using a Vanquish UPLC (Thermo Fisher Scientific, Waltham, MA) on a 100 x 2.1 mm Kinetix 1.7 µM C18 column (Phenomenex, Torrance, CA) using the following buffers. Buffer A: water (J.T.Baker, LC-MS grade) with 0.1% formic acid (Thermo Fisher Scientific, Optima LC/MS). Buffer B: acetonitrile (J.T.Baker, LC-MS grade) with 0.1% formic acid (Fisher Scientific, Optima LC/MS). Flow rate was set to 0.5 mL/min with a gradient of 0-1 min 5% B, 1-8 min 100% B, 8-10.9 min 100% B, 10.9-11 min 5% A, 11-12 min 5% B. Mass range was set to 100-1500 m/z, MS1 scan level resolution was set to 35K and MS2 scan resolution was set to 17.5K. Data is deposited in MassIVE (massive.ucsd.edu) under accession MSV000079760. The targeted dataset contained 21 samples, 1925 metabolic featues (33 of which were internal standards), with 6551(16.2%) missing data points (0, no detected signal).

#### Untargeted dataset

Dataset 2 was collected from NIST reference grade test materials (RGTM) containing homogenized fecal matter from subjects with vegan or omnivore diets (RGTM 10162, 10171, 10172, and 10173). Samples were collected from 18 individuals (9 vegan, 9 omnivore) with three technical replicates per individual. MS/MS data was acquired in positive mode on a Q Exactive Orbitrap (Thermo Fisher Scientific, Waltham, MA) using data-dependent acquisition. The untargeted dataset contained 54 samples, 42,44 metabolites, with 32,675 (14.3%) missing data points (0, no detected signal). Data is deposited in MassIVE (massive.ucsd.edu) under accession MSV000086989.

#### Data processing

Data processing was performed using the GNPS analysis ecosystem and MZmine3 (3.9.0)^46^. Raw data was first converted to .mzML using Proteowizard MSconvert (3.0.22287-170037b) before performing feature finding in MZMine3^47^. All feature finding parameters are included in Supplementary Files. Library annotation and molecular networking were performed using GNPS for Dataset 1 and can be accessed via the following link:

### Normalizations and Transformations

#### Total Ion Count Normalization

TIC normalization (TICN) was carried out by summation of all signals for each metabolite (*s*_*i*_) - Total Ion Count (TIC) - within a sample (*s*) and dividing each metabolite’s signal strength by the resulting sum.

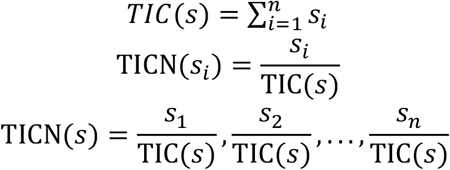

#### Centered Log Ratio Transformation

The centered log-ratio (CLR) transformation transforms compositional data into real number space by taking the logarithm of each sample’s component value and dividing by the geometric mean (GM) of all components, thereby eliminating the constant-sum constraint and making the data suitable for multivariate analysis. The generalized formula for the CLR transformation is shown below, where *s* is a sample vector, *s*_*i*_ is a component (metabolite signal) of the sample vector, CLR(*s*_*i*_) is an individual log ratio for a component of the vector, and CLR(*s*) is the transformed vector:

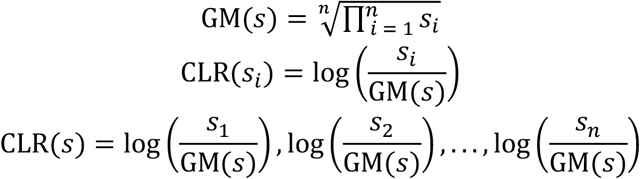

In the context of the metabolomics data sets, the raw metabolite signals are not inherently compositional. Before performing the CLR transformation for the data sets, each sample was first TIC-normalized to convert it into compositional form, then the CLR transformation was applied to each sample vector to yield a final TIC-CLR transformation. CLR calculations were performed using the decostand(x, method = “clr”, pseudocount = 1e-12) function within the vegan R library^48^.

#### Robust Centered Log Ratio Transformation

The robust centered log ratio (RCLR) transformation is an extension of the traditional CLR transformation which allows for the presence of zeros within datasets. The RCLR transformation ignores zero values and divides all non-zero values by the geometric mean of the observed features, followed by a log transformation on the non-zero log ratios; the zero values remain unchanged. After the transformation, the unchanged zero values must be handled. The RCLR transformation is calculated similarly to the CLR transformation, with the caveat that 0/missing signals are not included in the calculation:

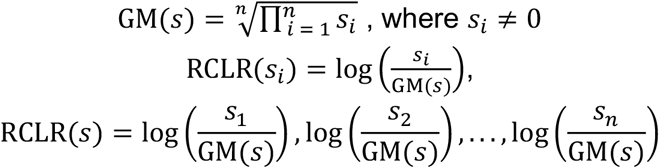

Similar to the CLR transformation, each sample was first TIC-normalized to convert it into compositional form, then the RCLR transformation was applied to each sample vector to yield a final TIC-RCLR transformation. RCLR calculations were performed using the decostand(x, method = “rclr”) function within the vegan R library.

#### Proof that compositional transformation on TIC-normalized data removes compositionality

Let the vector of data 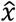 we consider be defined as the following

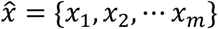

We say that 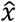 is a vector in ℝ^*m*^ and denote the *i*-th entry of 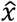 as *x*_*i*_. We define the TIC-normalized data vector 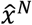 by its *i*-th entry as:

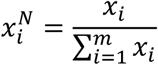

Each entry is the corresponding entry in the non-normalized vector divided by the sum of all entries. This normalization can be shown to provide compositional data because the sum of the elements of the normalized vector, 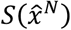, can be written as:

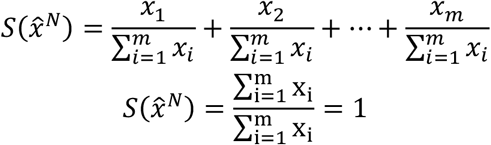

We define the CLR transformation of the TIC-normalized vector by its *i*-th entry as:

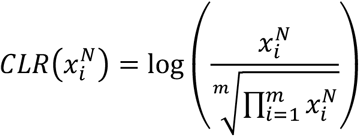

Similarly, the CLR transformation of the un-normalized data vector by its *i*-th entry is defined as:

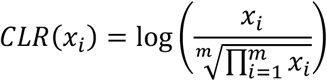

We can expand the CLR transform of the normalized vector as the following:

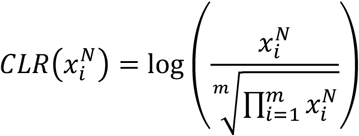

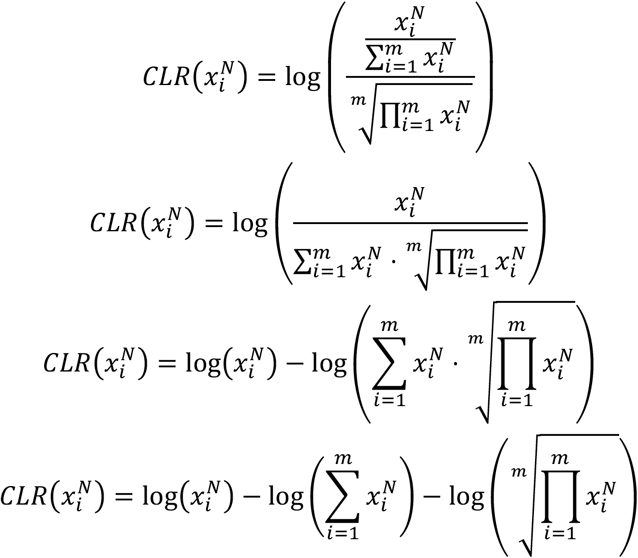

A similar expansion results in the *i*-th entry of the CLR transform of the un-normalized vector being written as the following:

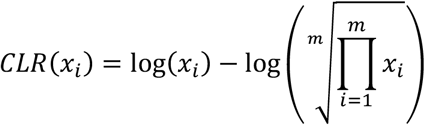

We can see that the two different treatments of the data vector only differ by the constant term 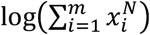 and are notably similar in terms of interpretation as noted in the main body text.

We then observe the sum of the TIC-CLR transformed data vector to determine if the if the compositionality has been removed:

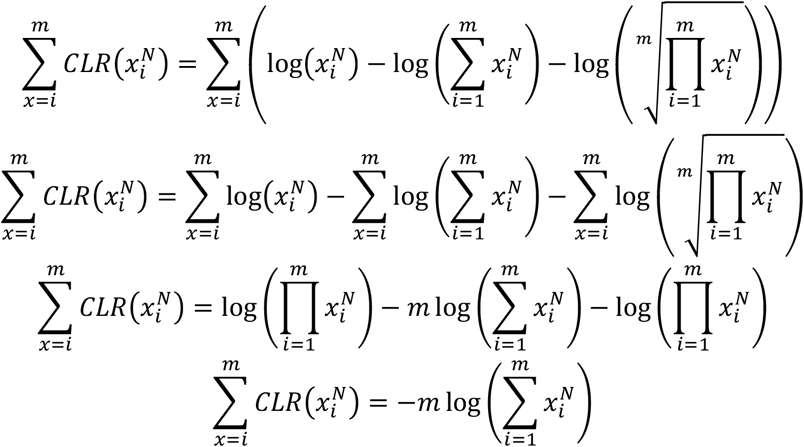

We find that the resulting sum is equal to the negative logarithm of the sum of entries of the original vector multiplied by the size of the vector, thereby showing that the entries do not sum to a fixed value that is not dependent on the data and thus, the data lacks compositionality.

#### Minimum Value Imputations

Minimum value imputations were carried out for RCLR transformations, substituting non-transformed 0/missing signal values with the minimum value of TIC-RCLR transformed feature table.

#### k-Nearest Neighbors Imputations

The k-Nearest Neighbors imputations were carried out with the R package VIM^49^ using the kNN function. The kNN function uses the Gower Distance to calculate similarity between samples/variables as the method allows for mixed data types. In the case of purely quantitative data types (used in this study), the Gower Distance calculations only employ Manhattan Distance. The kNN method will raise an error if any features (metabolites) contain entirely missing values. The method arguments used were k = 3 (three nearest neighbors), numFun = mean (averages the value of the 3 nearest neighbors), useImputedDist = FALSE, and default values for all other arguments.

#### Random Forests Imputations

The Random Forests imputations were generated using the R package missForest^50^, with the self-named function missForest(). The parameters used were 10 for *maxiter* (maximum number of iterations if convergence has not yet been reached), 100 for *ntree* (number of trees to grow in each forest), and default values for all other arguments.

The general steps to the iterative algorithm are (imputations are made in order for samples/variables with the least amount of missing values, moving towards samples/variables with more missing values):

1. Initialize the imputation process by making initial guesses for missing values with the mean of the matrix
2. For each sample/variable with missing values, a random forest is trained using other samples/variables as predictors; the training is done only on observed values.
3. The trained random forests model predicts the missing values for the sample/variable
4. The missing values for the sample/variable are updated with each iteration.
5. The process is repeated iteratively until the imputed values reach convergence (or a predefined number of iterations is specified).
6. After convergence (or maximum iterations), the imputed data set is returned.

Details of the full algorithm are specified in the source publication.

### Targeted Metabolomics Dataset Methods

To assess the effects of machine learning imputations and model-based imputation when the missing mechanism is MNAR (data is missing systematically at the lowest concentration/at the limit of detection), the targeted dataset was utilized. This dataset contained 41 internal standard metabolites which were spiked-in at fixed concentrations of 10 pM, 100 pM, 1 nM, 10 nM, and 100 nM. For machine learning imputations, all missing values were imputed with both kNN and RF with the parameters discussed above. For modeling-based imputations, a logistic regression was fit to the detected data, then used to predict values for missing data.

### Untargeted Metabolomics Dataset Methods

The effectiveness of machine learning based imputations for MCAR-type and MNAR-type data was assessed using the untargeted dataset. To directly compare the performance of RF and kNN imputations, the dataset was TIC-RCLR transformed, then missing values were randomly assigned at 20%, 40%, 60%, and 80% intervals using the prodNA() function within the missForest package. Each dataset with missing values at the different intervals was imputed separately, then imputation performance was assessed by normalized root mean square error (NRMSE) with the missForest function nrmse(), which accepts the imputed dataset, the dataset with missing (NA) values, and the true dataset (TIC-RCLR transformed, no missing/imputed values) as arguments. Metabolite-level imputation accuracy was also assessed by absolute distance between true value and imputed value at three discrete intervals: metabolites which had <=10% detected signals across all samples, metabolites with 50% detected signals across all samples, and metabolites which had >=90% detected signals across all samples. Correlation was determined with cor.test(method = “pearson”).

## ACKNOWLEDGEMENTS

Dennis D. Krutkin’s graduate studies are supported by a Division of Research and Innovation Doctoral Fellowship.

## DATA AND CODE AVAILABILITY

The targeted metabolomics data set used for analysis is deposited in MassIVE (massive.ucsd.edu) under accession MSV000079760. The untargeted metabolomics data set used for analysis is deposited in MassIVE under accession MSV000086989. The R Markdown scripts used for analyses and visualizations are available at https://github.com/ddkrutkin/metabolomics_imputations

