## Supplemental Figures for "To impute or not to impute in untargeted metabolomics - that is the compositional question"

### Supplementary Figures (CLR/RCLR on Raw Data)

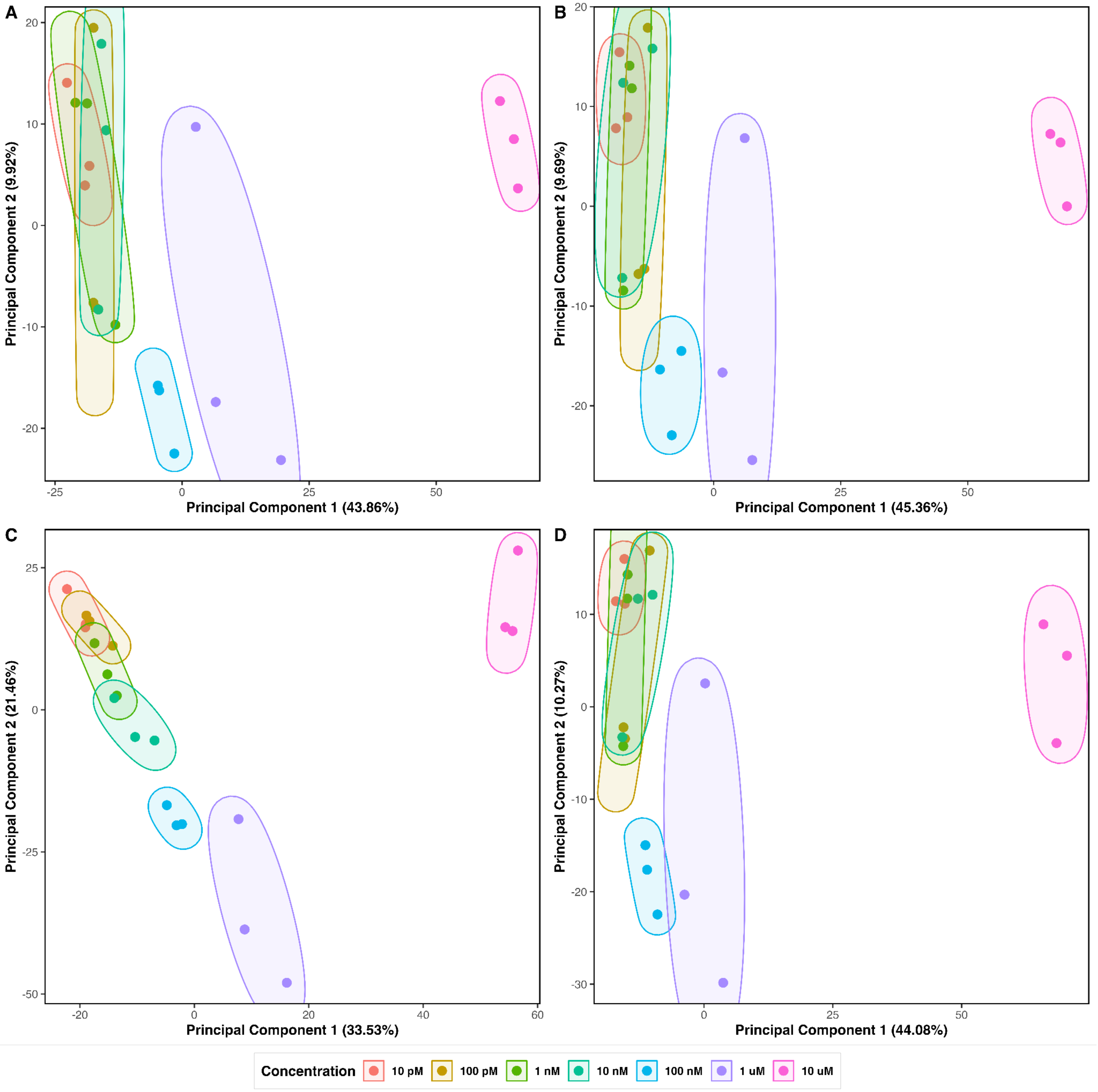


**Supplemental Figure 1.** PCA plots for different transformations on targeted metabolomics data; **A**. PCA plot for raw data, **B**. PCA plot for TIC normalized data, **C**. PCA plot for CLR transformed data (pseudocount = 1e-12), **D**. PCA plot for RCLR transformed data, followed by minimum value imputation.

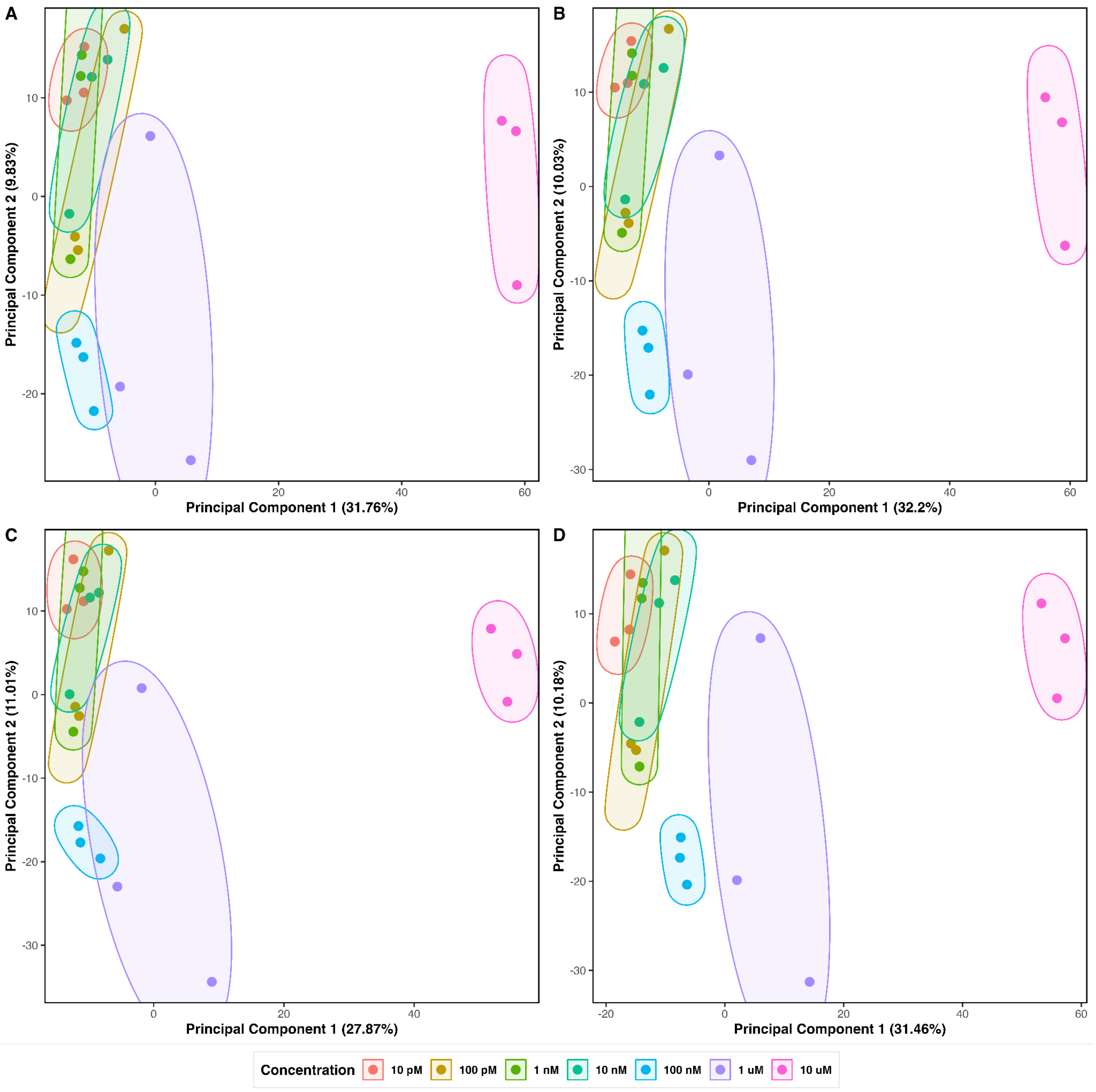


**Supplemental Figure 2.** PCA plots of the targeted metabolomics data using machine learning imputations and two different log-ratio transformation methods. **A**. RCLR transformed raw data, followed by RF imputation, **B**. RCLR transformed raw data, followed by kNN imputation, **C**. RF imputation on raw data, followed by CLR transformation, **D**. kNN imputation on raw data, followed by CLR transformation.

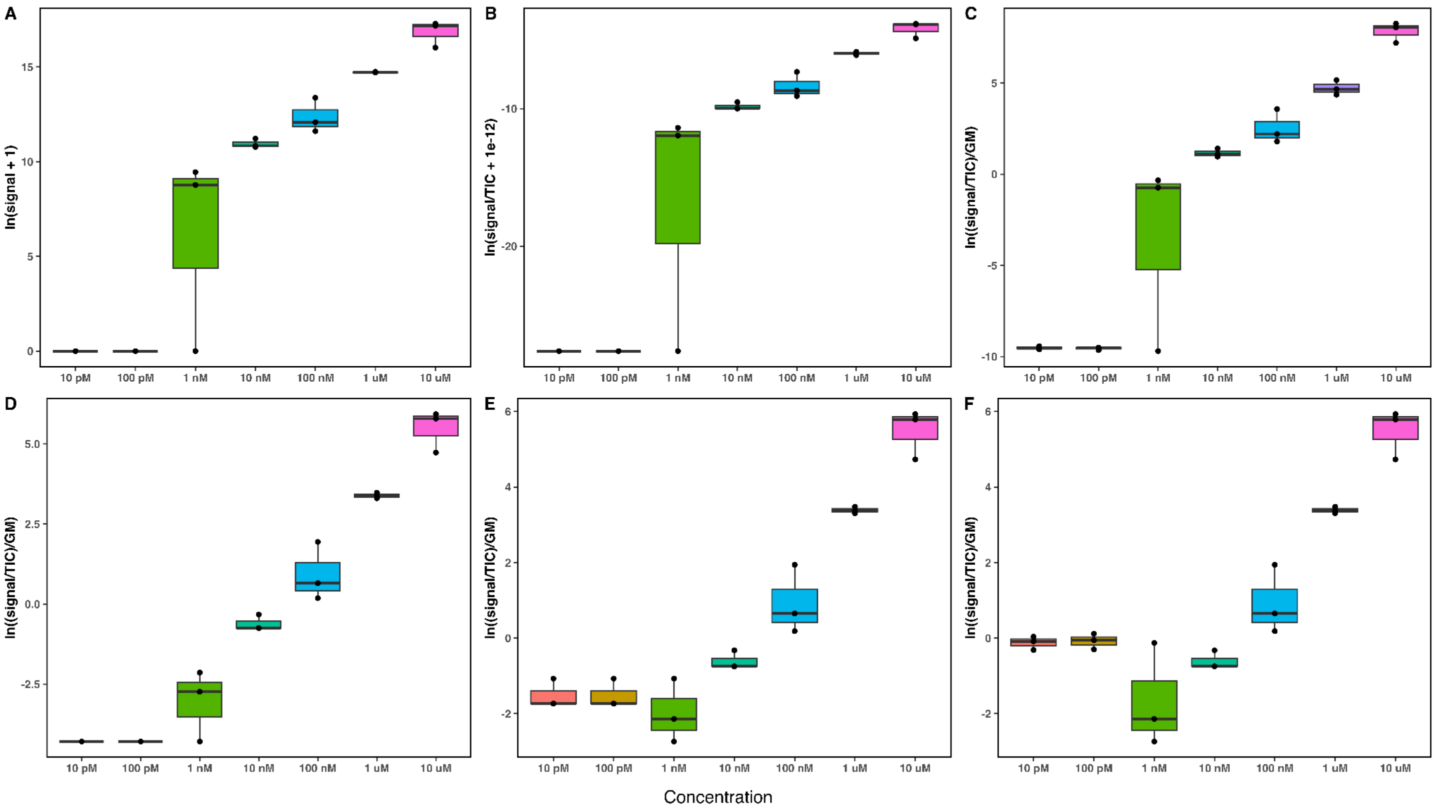


**Supplemental Figure 3.** Boxplots of transformations and imputations for aspartame, where the 2 lowest concentration thresholds have all missing data; **A**. Raw data plotted on a natural log scale, **B**. TIC normalized data plotted on a natural log scale, **C**. CLR transformed raw data (pseudocount = 1), **D**. RCLR transformed raw data, followed by minimum value imputation, **E**. RCLR transformed raw data, followed by kNN imputation, **F**. RCLR transformed raw data, followed by RF imputation.

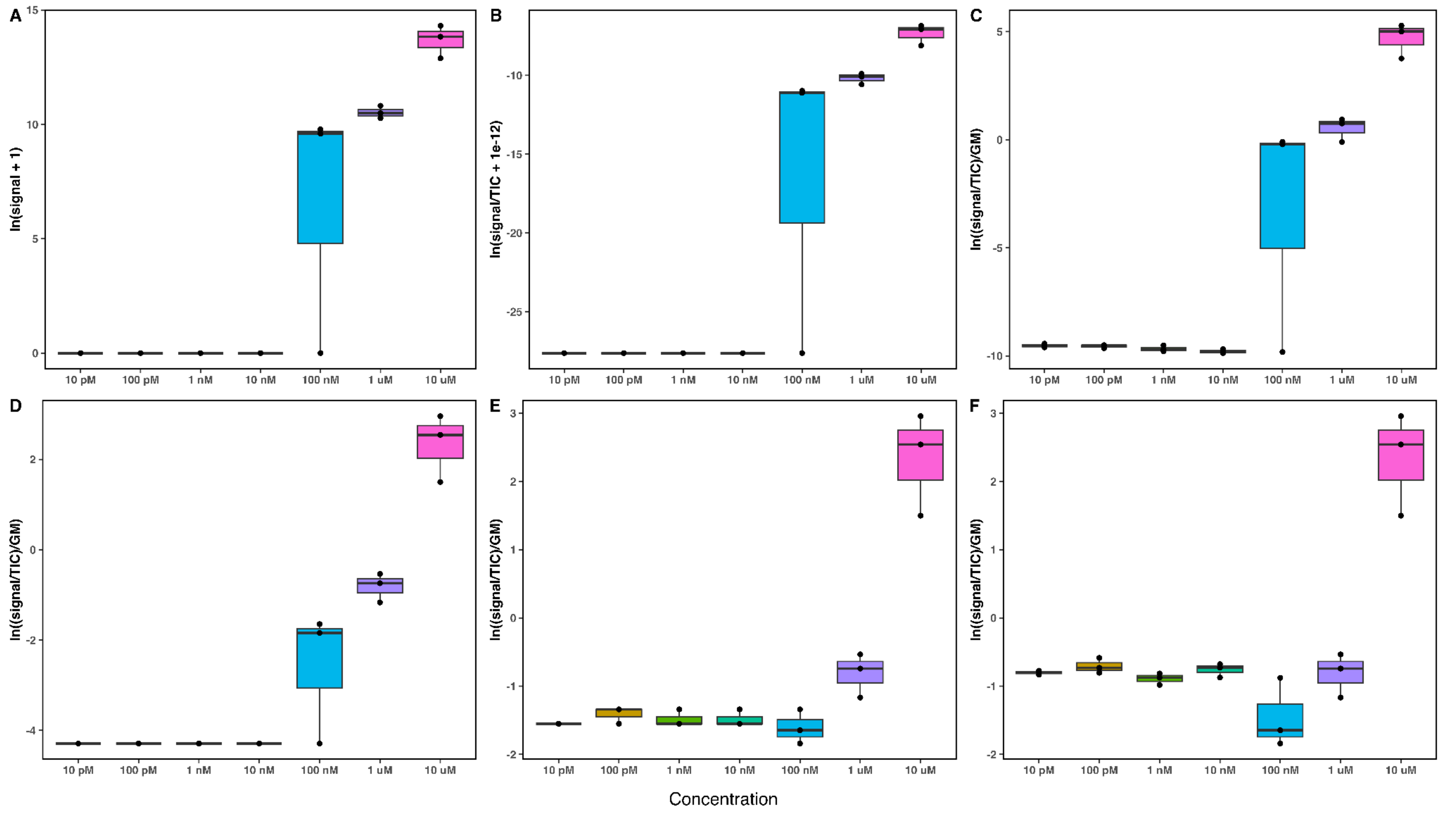


**Supplemental Figure 4.** Boxplots of transformations and imputations for estrone, which is missing signals for the four lowest concentrations; **A**. Raw data plotted on a natural log scale, **B**. TIC normalized raw data plotted on a natural log scale, **C**. CLR transformed raw data (pseudocount = 1), **D**. RCLR transformed raw data, followed by minimum value imputation, **E**. RCLR transformed raw data, followed by kNN imputation, **F**. RCLR transformed raw data, followed by RF imputation

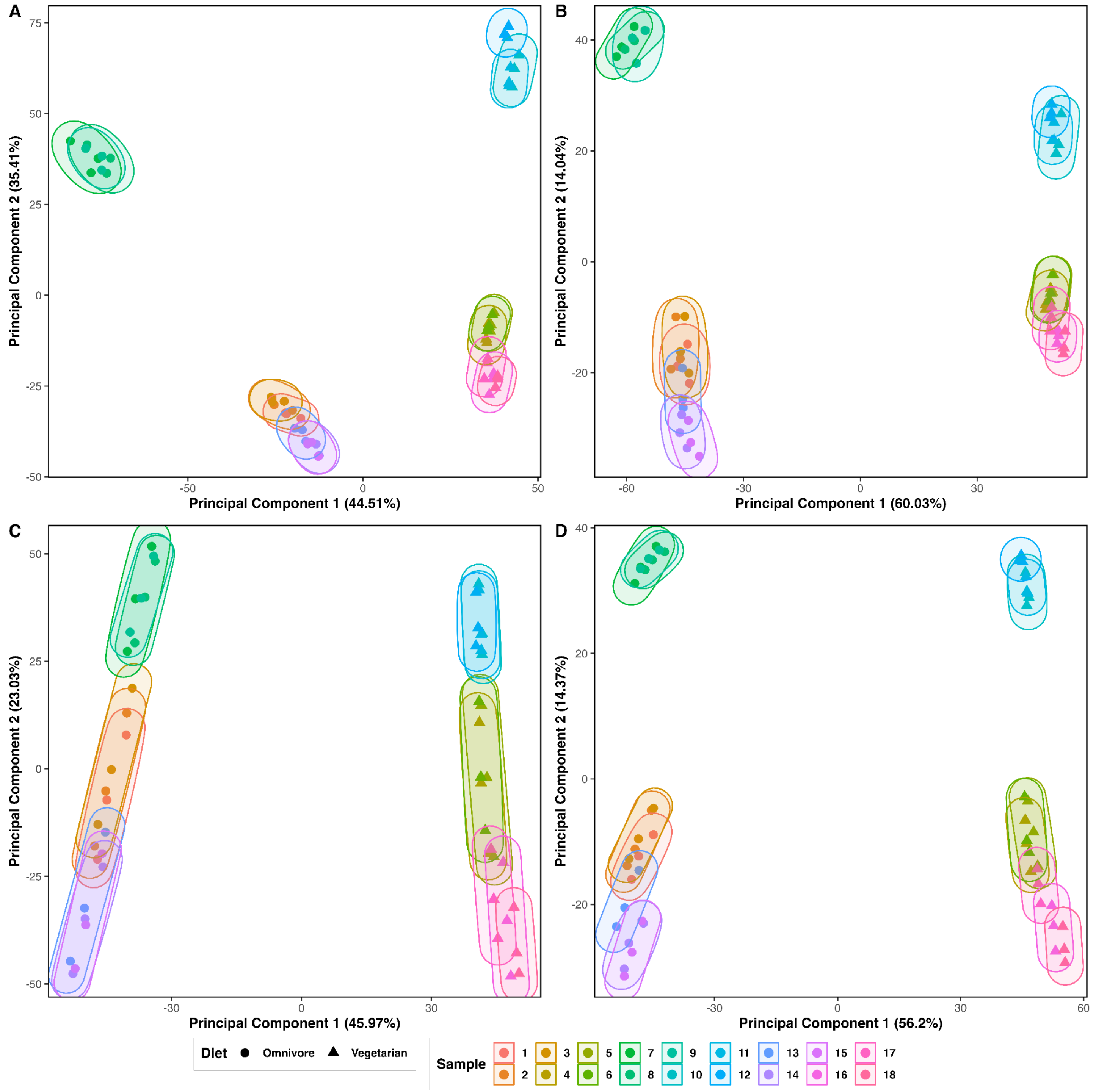


**Supplemental Figure 5.** PCA plots for different transformations on untargeted metabolomics data; **A**. PCA plot for raw data, **B**. PCA plot for TIC normalized data, **C**. PCA plot for CLR transformed data (pseudocount = 1), **D**. PCA plot for RCLR transformed data, followed by minimum value imputation.

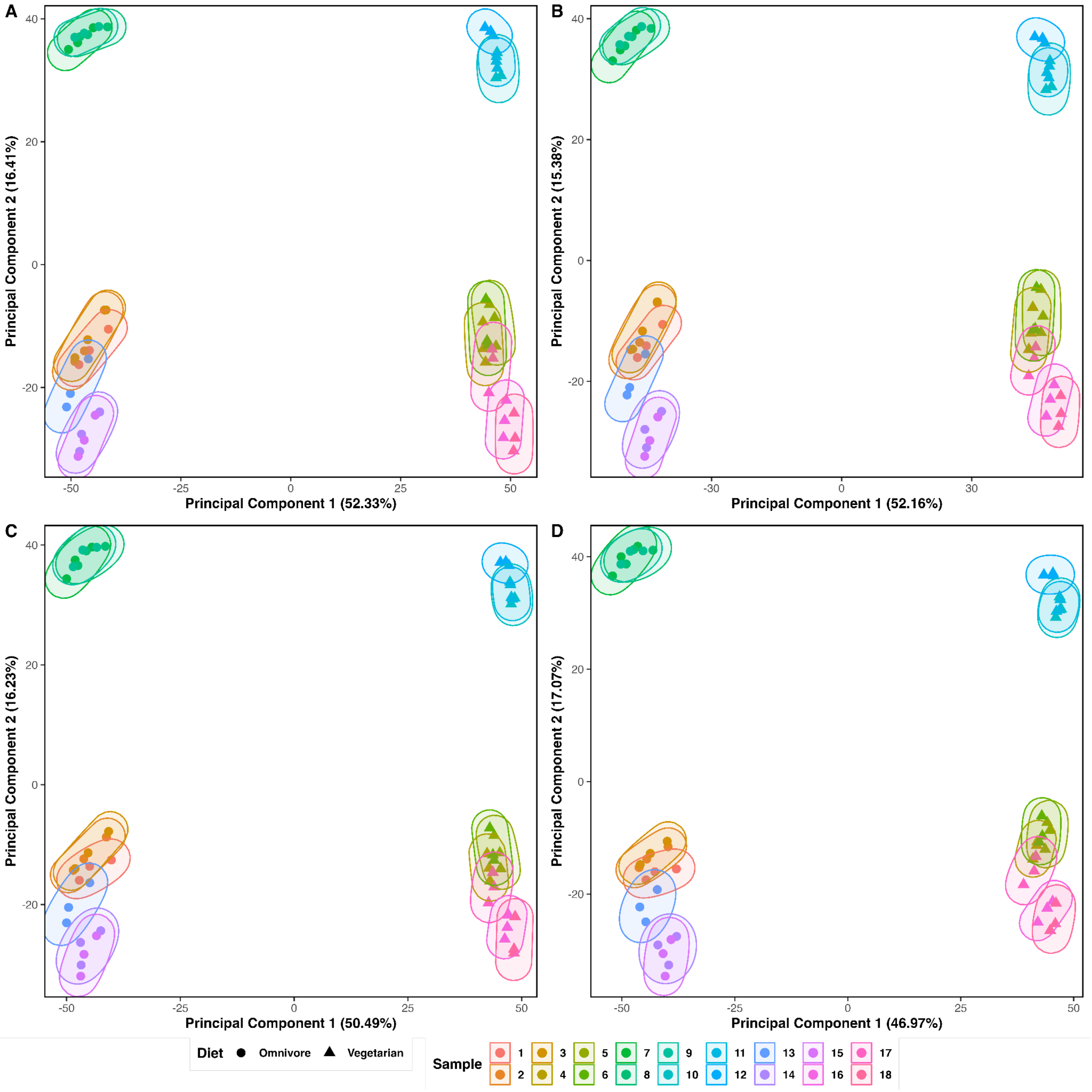


**Supplemental Figure 6.** PCA plots of the targeted metabolomics data using machine learning imputations and two different log-ratio transformation methods. **A**. RCLR transformed raw data, followed by RF imputation, **B**. RCLR transformed raw data, followed by kNN imputation, **C**. RF imputation on raw data, followed by CLR transformation, **D**. kNN imputation on raw data, followed by CLR transformation.

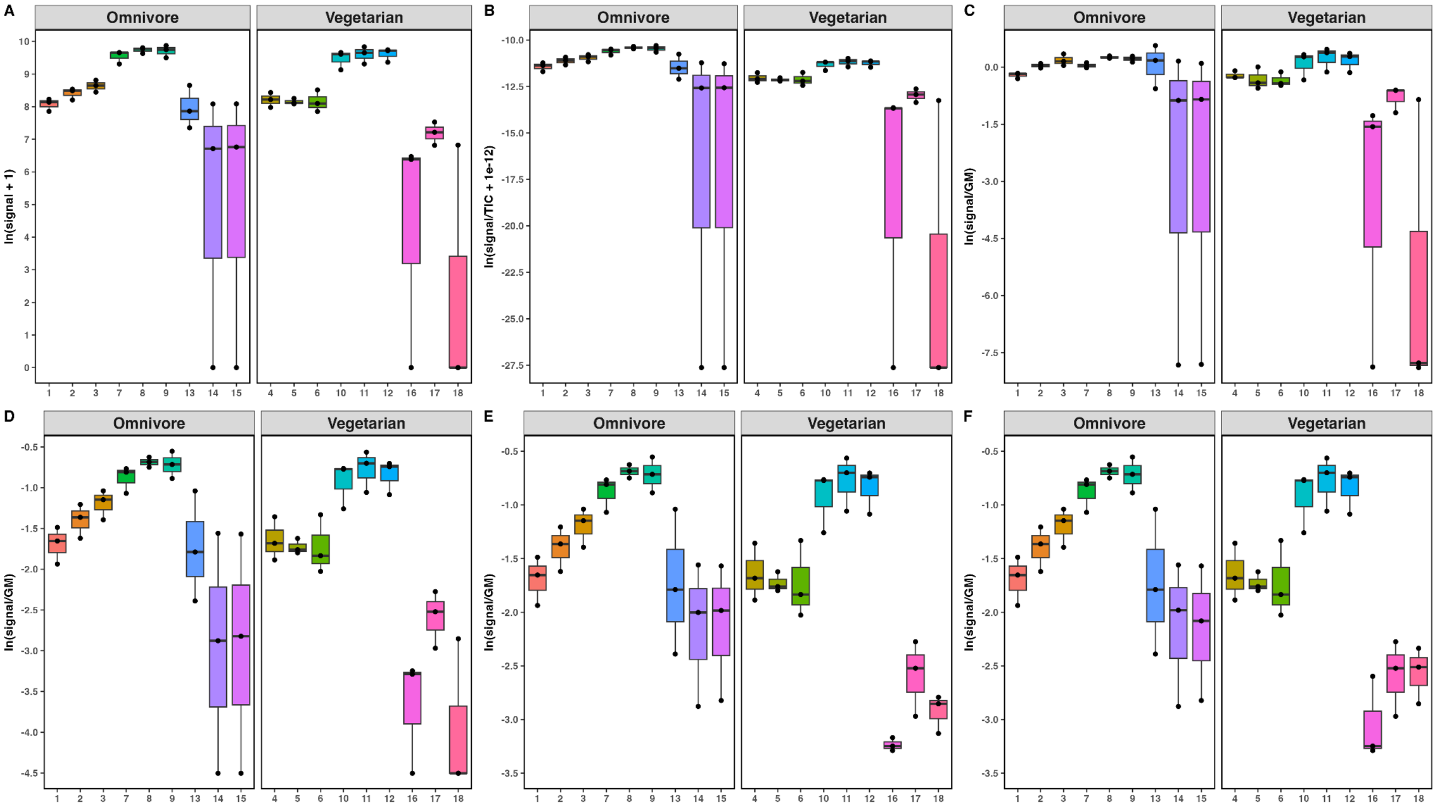


**Supplemental Figure 7**. Boxplots of transformations and imputations for an unknown metabolite with >90% detected signals across all samples in the untargeted dataset; **A**. Raw data plotted on a natural log scale, **B**. TIC normalized data plotted on a natural log scale, **C**. CLR transformed data (pseudocount = 1), **D**. RCLR transformed data, followed by minimum value imputation, **E**. RCLR transformed data, followed by kNN imputation, **F**. RCLR transformed data, followed by RF imputation.

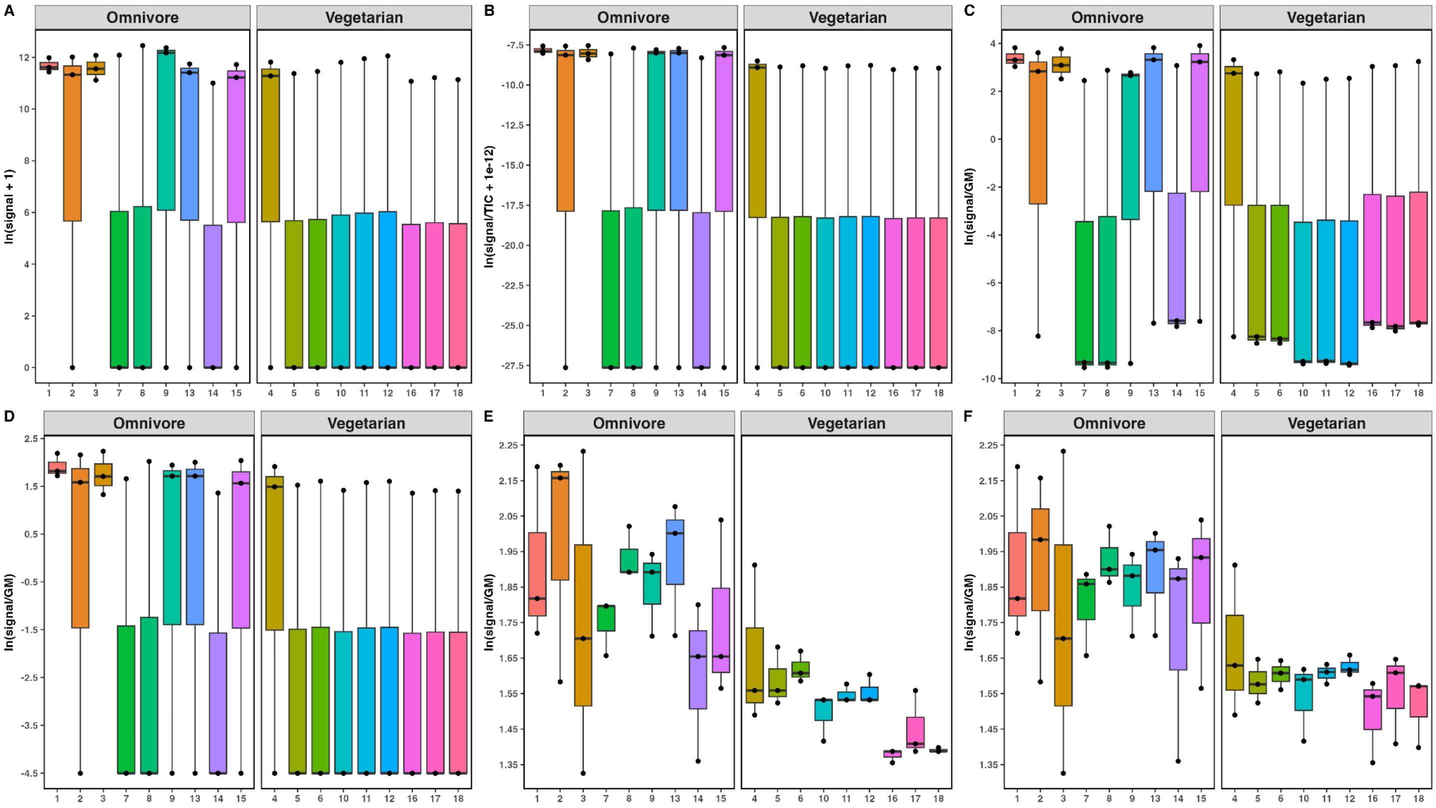


**Supplemental Figure 8**. Boxplots of transformations and imputations for an unknown metabolite with 50% detected signals and 50% missing signals (where the data is MCAR) across all samples in the untargeted dataset; **A**. Raw data plotted on a natural log scale, **B**. TIC normalized data plotted on a natural log scale **C**. CLR transformed data (pseudocount = 1), **D**. RCLR transformed data, followed by minimum value imputation, **E**. RCLR transformed data, followed by kNN imputation, **F**. RCLR transformed data, followed by RF imputation.

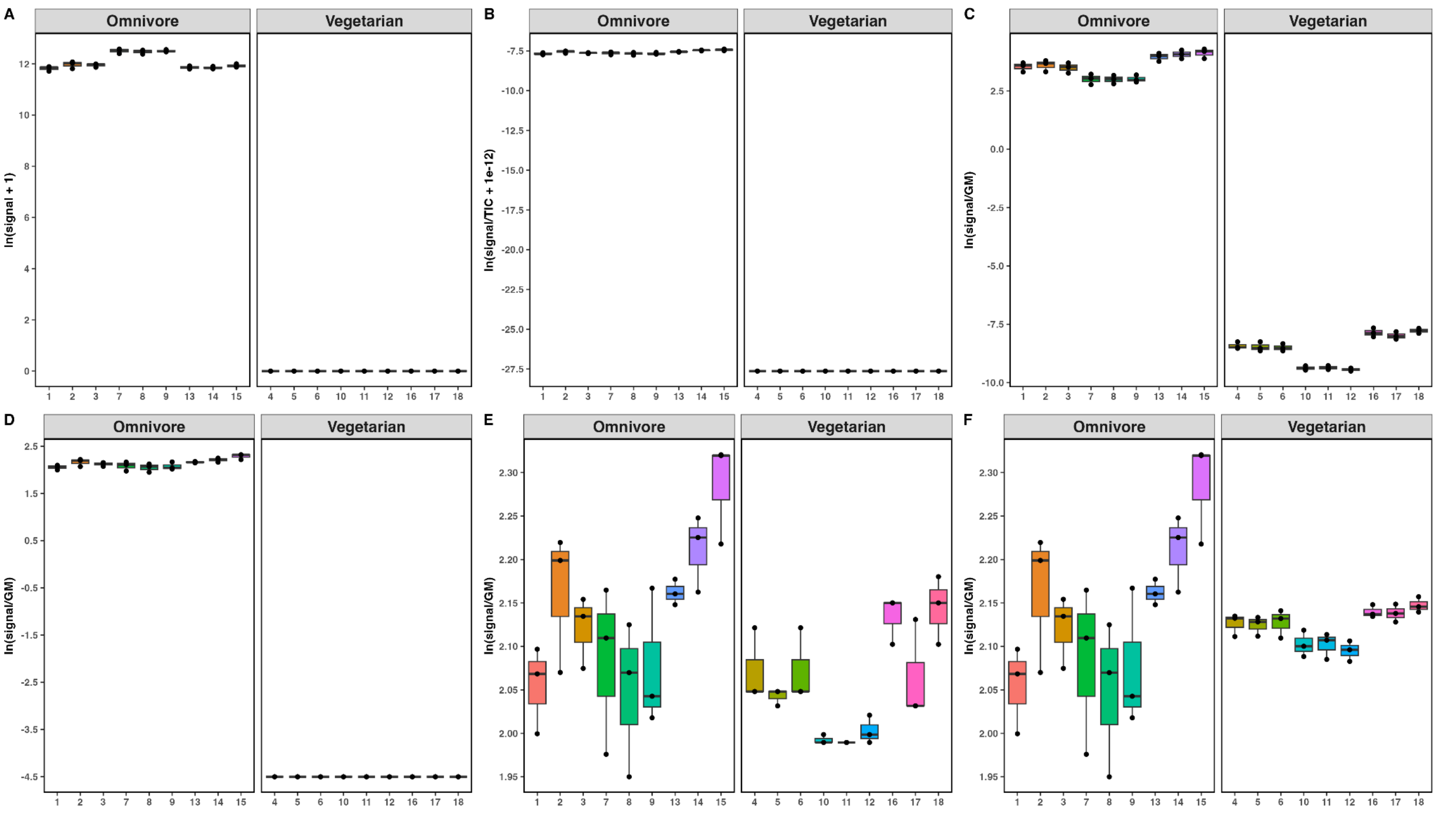

**Supplemental Figure 9**. Boxplots of transformations and imputations for an unknown metabolite with 50% detected signals and 50% missing signals (where the data is MNAR) across all samples in the untargeted dataset; **A**. Raw data plotted on a natural log scale, **B**. TIC normalized data plotted on a natural log scale, **C**. CLR transformed data (pseudocount = 1), **D**. RCLR transformed data, followed by minimum value imputation, **E**. RCLR transformed data, followed by kNN imputation, **F**. RCLR transformed data, followed by RF imputation.

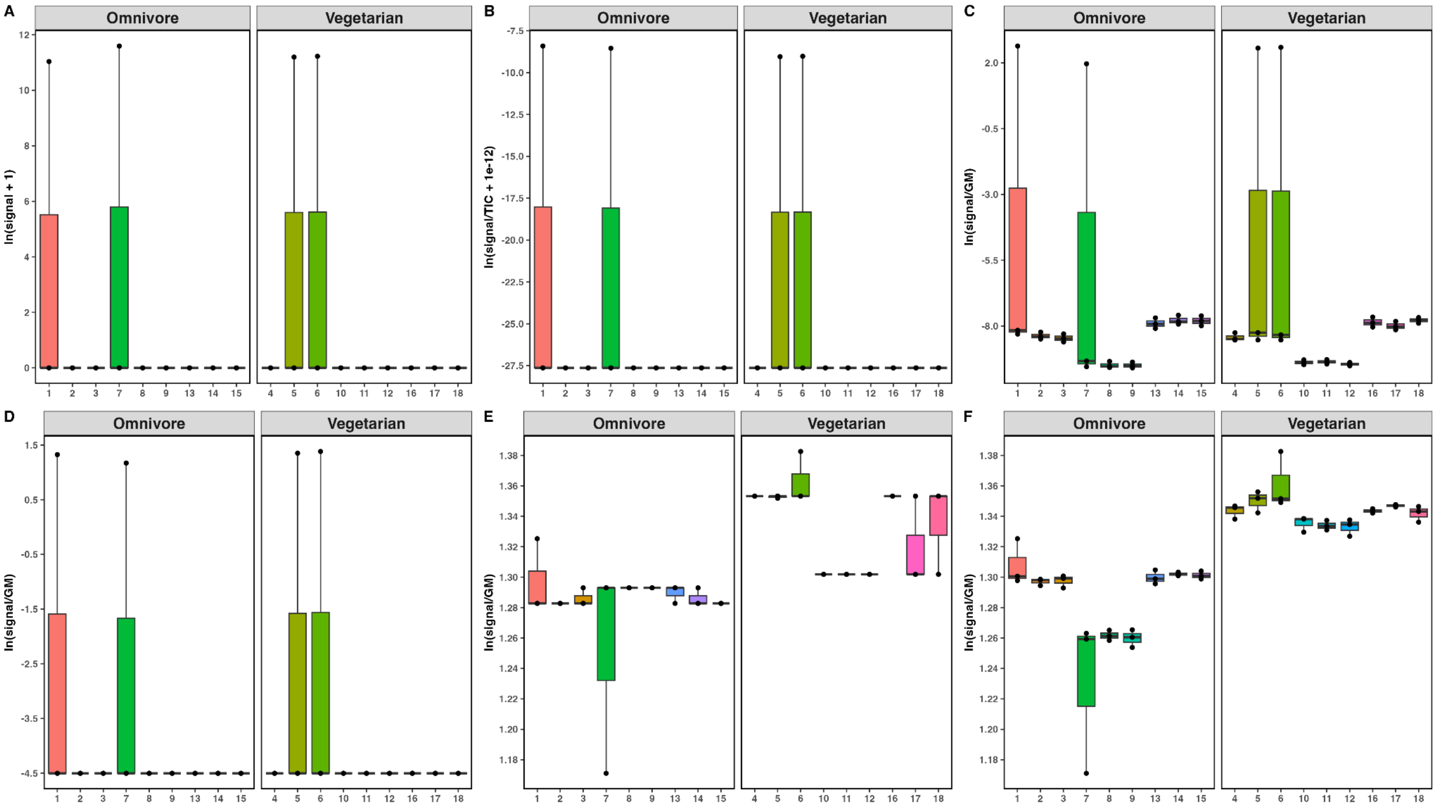


**Supplemental Figure 10**. Boxplots of transformations and imputations for a metabolite <10% detected signals across all samples in the untargeted dataset. **A**. Raw data plotted on a natural log scale, **B**. TIC normalized data plotted on a natural log scale, **C**. CLR transformed data (pseudocount = 1), **D**. RCLR transformed data, followed by minimum value imputation, **E**. RCLR transformed data, followed by kNN imputation, **F**. RCLR transformed data, followed by RF imputation.
